# Co-opted cytosolic host proteins form unique condensate substructures within the membranous tombusviral replication organelles

**DOI:** 10.1101/2023.07.26.550743

**Authors:** Wenwu Lin, Peter D. Nagy

**Affiliations:** Department of Plant Pathology, University of Kentucky, Lexington, United States of America

## Abstract

Positive-strand RNA viruses co-opt intracellular organellar membranes for the biogenesis of viral replication organelles (VROs). The membranous VROs are the sites of viral replication. Tombusviruses co-opt numerous host proteins, many of them cytosolic, not membrane-bound, such as the glycolytic and fermentation enzymes. It is currently not known what type of molecular organization keeps these enzymes enriched and sequestered within the membranous VROs. Here, we show evidence that the tomato bushy stunt virus (TBSV) p33 and the closely-related carnation Italian ringspot virus (CIRV) p36 replication proteins sequester several co-opted cytosolic proteins in unique condensate substructures associated with the membranous VROs. We find that TBSV p33 and CIRV p36 replication proteins form droplets *in vitro* organized by their intrinsically disordered region (IDR). The replication proteins also organize the partitioning of several co-opted host proteins to p33/p36 droplets. We also show that the VRO-associated condensates containing co-opted glycolytic and fermentation enzymes are critical for local ATP production within the VRO to support the energy need of virus replication. A short segment with charged amino acids in p33/p36 IDR plays a critical role in droplet formation *in vitro*, in organizing the VRO-associated condensates and affects both ATP production in VROs and viral replication. We find that the co-opted ER membranes and actin filaments form meshworks within and around the condensate, likely contributing to the unique composition and structure of VROs. We propose that the p33/p36 replication proteins organize liquid-liquid phase separation of co-opted highly concentrated host proteins in condensate substructures within VROs to allow for the dynamic regulation of viral replication. Overall, we demonstrate that distinct substructures co-exist in tombusvirus VROs. These are the subverted membranes and condensate substructures, both of which are critical for VRO functions. The p33/p36 replication proteins induce and connect the two substructures within the VROs. Interfering with condensate formation in VROs might open up new antiviral approaches.

## Introduction

Positive-strand (+)RNA viruses replicate on subverted subcellular membranes by forming unique viral replication organelles (VROs) [1–5]. The VROs are remodeled and deformed cellular membranes, which contain membrane-bound viral replicase complexes consisting of viral replication proteins, viral RNAs and co-opted host proteins and represent the sites of viral replication [1, 5–11]. The VROs concentrate all replication factors into a distinct cytosolic area for efficient RNA synthesis and spatial and temporal organization of viral replication. The VROs also protect the fragile viral RNAs from degradation by host ribonucleases and hinder the recognition of viral components by the host antiviral defences [12–17]. Overall, assembly of VROs is a key step during replication of (+)RNA viruses in the infected cells. In addition to the subverted subcellular membranes, (+)RNA viruses have to co-opt numerous cytosolic host proteins and recruit them to VROs for providing key pro-viral functions. Yet, the robust recruitment and enrichment process of the cytosolic proteins into membranous VROs is currently poorly understood.

Tomato bushy stunt virus (TBSV), a plant-infecting tombusvirus, serves as an outstanding model to dissect virus-induced complex rearrangements of cellular membranes and alterations in lipid and other metabolic processes during infections [18–20]. Tombusviruses belong to the large Flavivirus-like supergroup that includes important human, animal and plant pathogens. TBSV has a ∼4.8 kb single component (+)RNA genome. Among the five proteins coded by the TBSV genome, there are two essential replication proteins. These are the abundant p33 and p92^pol^, the latter of which is the RdRp protein. P92^pol^ is translated from the genomic RNA via readthrough of the translational stop codon in p33 ORF [21]. P33 replication protein is an RNA chaperone and selects and recruits viral (+)RNA for replication [21–23]. The emerging picture is that p33 is the master regulator of VRO biogenesis [3, 24, 25]. Tombusviruses hijack various cellular compartments and pathways for VRO biogenesis [24–26]. These include peroxisomes by TBSV or mitochondria by the closely-related carnation Italian ringspot virus (CIRV). TBSV VRO consists of clustered peroxisomes (5-to-20) with vastly modified lipid compositions, which bear the numerous invaginated spherules, the sites of viral RNA synthesis [16, 18]. Subdomains of the ER membranes are also co-opted into VROs to form extensive membrane contact sites with the co-opted peroxisomes for sterol enrichment in VROs [18, 27–30]. Moreover, Rab1- and Rab5-positive vesicles and the retromer tubular transport carriers are also subverted to supply additional membrane surfaces and host factors for VRO biogenesis [24, 31–33]. Importantly, numerous cytosolic host factors are also co-opted, which are not membrane bound. These include several co-opted glycolytic and fermentation enzymes to generate ATP locally within the VROs [34–37]. Interestingly, TBSV p33 replication protein targets the proteasomal Rpn11 key protein interaction ‘hub’, which is co-opted to facilitate the recruitment of glycolytic and fermentation enzymes into VROs [38]. It is currently not known how all these co-opted cytosolic proteins are sequestered and enriched within VROs.

A recently emerging concept in cell biology is that the cytosol is not uniform, but organized into intracellular assemblies, called biomolecular condensates under various conditions [39–41]. Formation of condensates provides a mechanism of concentrating and segregating a set of cellular components in a spatially defined compartment for various biological activities. The formation of condensates is driven by liquid-liquid phase separation, which is induced by scaffolding proteins or nucleic acids. During phase separation, proteins form interconnected networks through multivalent, low-affinity binding. The transient condensates can undergo fusion or fission and exchange components with the surrounding cytosol. Phase separation can also lead to gel-like condensates, which are more stable assemblies [42–44]. Condensates could also associate with subcellular membranes and lipids [45, 46]. Altogether, phase separation has emerged as one of the major principles of subcellular organization.

In this paper, we show evidence that the TBSV p33 and the CIRV p36 replication proteins induce phase separation sequestering several co-opted host proteins in condensate substructures, which are critical for VRO functions in plant cells. The VRO-associated condensate represents a distinct phase from the membranous substructures, the latter of which contain the viral replicase complexes. We also show that the VRO-associated condensate is required for local ATP production by the co-opted glycolytic enzymes. We propose that the co-existence of the membranous and the condensate substructures within VROs allows the dynamic regulation of viral replication.

## Results

### Co-opted host proteins are present in biomolecular condensates in the membranous tombusvirus replication organelles

Many co-opted host proteins are sequestered and enriched in the membranous tombusvirus replication organelles (VROs) [3, 47]. However, most of these co-opted host proteins are not membrane bound. What makes these proteins trapped in the VROs? For example, the co-opted glycolytic and fermentation enzymes are soluble proteins, but stably and tightly associated with VROs during tombusvirus infections [34, 48]. It is currently not known what type of molecular organization/structure keeps these enzymes sequestered within the membranous VROs. We have tested if formation of condensate substructures might contribute to VRO biogenesis in *Nicotiana benthamiana* host. Indeed, fluorescence recovery after photobleaching (FRAP) experiments in plant cells showed that the co-opted fluorescent Fructose-1,6-bisphosphate aldolase tagged with Red fluorescent protein (RFP-Fba2) glycolytic enzyme moved into the photobleached area within the TBSV p33 or CIRV p36 replication proteins-induced VROs (Fig 1A-C), indicating condensate structure, which allows internal movement of proteins. The fluorescence signal recovery was 50-to-60% and relatively slow (2 min), suggesting that Fba2 is present in a gel-like condensate within VROs induced by TBSV p33 or CIRV p36 replication proteins. The membrane-bound TBSV p33 or CIRV p36 fluorescence signals were not recovered, because the co-opted peroxisomal or mitochondrial membranes restrict the movement of these replication proteins (Fig 1A-C). We also tested the co-opted glycolytic Glyceraldehyde-3-phosphate dehydrogenase (GAPDH or GAPC), Phosphoglycerate kinase (Pgk1) and Pyruvate kinase (PK) and the fermetation enzymes Pyruvate decarboxylase (Pdc1) and Alcohol dehydrogenase (Adh1) in the FRAP assay. The fluorescence signals for all these co-opted host proteins were recovered 30-to-70% within the TBSV p33 or CIRV p36-induced VROs in 2 min (Fig S1&S2), indicating that all these proteins were present in the VRO-associated condensates. The fluorescence signals for a few co-opted glycolytic proteins, such as GAPDH, PK and Pdc1, were lower at 2 min after photobleaching, but gradually recovered up to the 10 min time point (Fig S2). In the control FRAP experiment, the fluorescence recovery of RFP-Adh1 in the cytosol, in the absence of TBSV, was rapid and complete (Fig S1C), suggesting that Adh1 does not form condensate under these experimental conditions.

**Fig 1.**
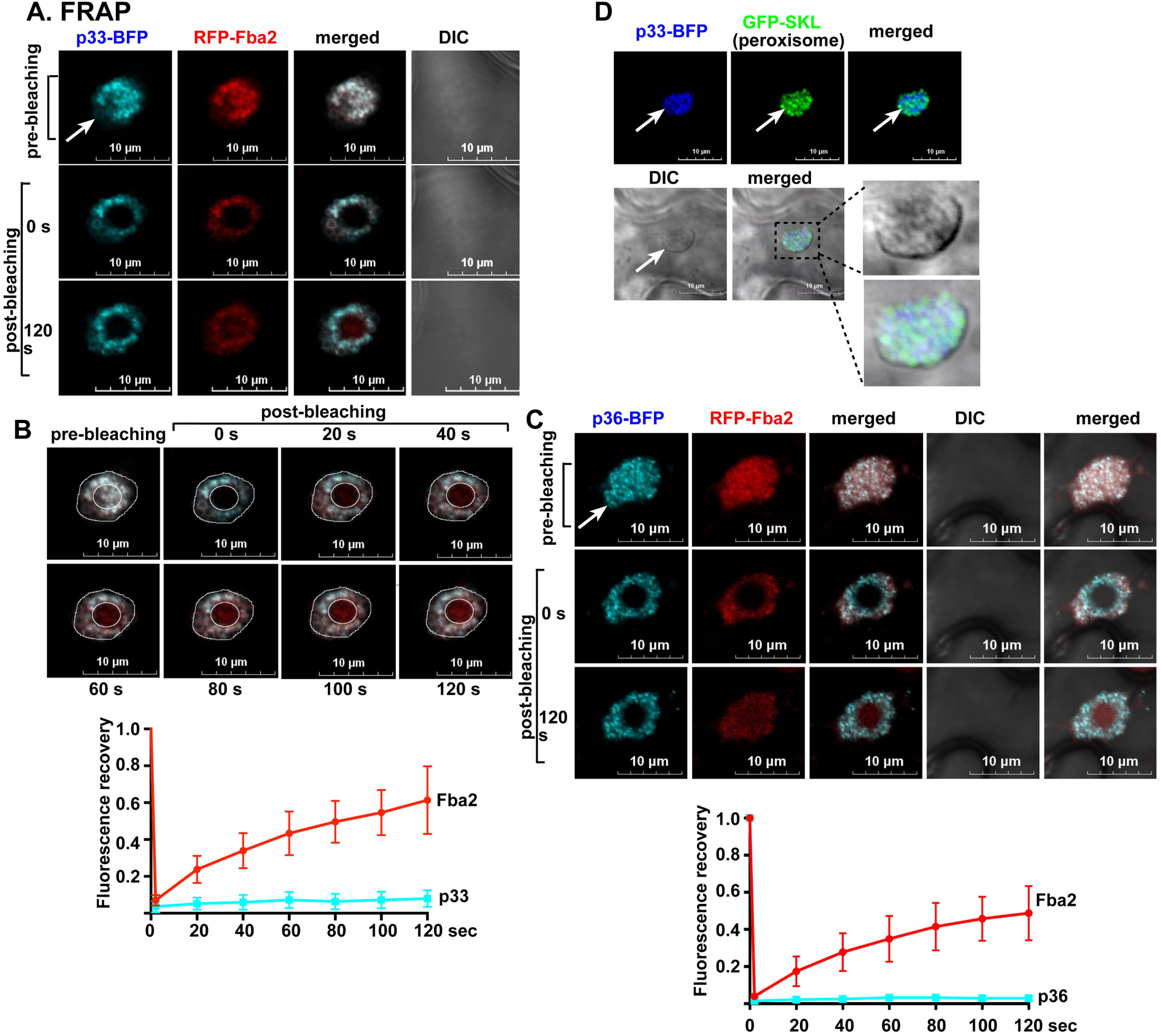
The co-opted Fba2 glycolytic enzyme is present in condensate substructure in tombusvirus VROs. (A) FRAP analysis shows the partial fluorescence recovery of RFP-Fba2 signal after photobleaching within a single VRO induced by TBSV p33-BFP in a *N. benthamiana* cell. An arrow points at the large VRO consisting of clustered peroxisomes. Note that the membrane-bound p33-BFP signal was not recovered. Scale bars represent 10 μm. (B) Time course analysis of fluorescence recovery of the RFP-Fba2 signal after photobleaching is shown within a single VRO. The merged images of p33-BFP and RFP-Fba2 are presented to show the extent of fluorescence recovery. 6-to-8 VROs from three independent plants were tested. (C) FRAP analysis shows the fluorescence recovery of RFP-Fba2 signal after photobleaching within a single VRO induced by CIRV p36-BFP in a *N. benthamiana* cell. The graph shows the extent of fluorescence recovery. (D) Confocal and DIC images show the VRO (pointed at by an arrow) induced by the TBSV p33-BFP.

We confirmed that GAPDH, PK and Fba2 showed partial fluorescence recovery in TBSV or CIRV-infected *N. benthamiana* cells using FRAP assay (Fig S3). As expected, fluorescence signals for the peroxisomal luminal marker (GFP-SKL) or the mitochondrial marker (GFP-Tim21) did not recover, because the organellar limiting membranes restrict the movement of peroxisomal or mitochondrial marker proteins. All these data indicate that the co-opted glycolytic and fermentation enzymes are enriched and sequestered in a VRO-associated condensate, which constitutes a new distinct substructure within the membranous VROs of tombusviruses. We call this new substructure as virus-induced replication condensate, or vir-condensate, which is a dynamic portion of the VROs. DIC images showed that the VROs are separated from the cytosol by a dense layer (Fig 1D), which may represent the highly concentrated co-opted molecules in vir-condensates.

Not only condensate formation, but other mechanism might also lead to substantial enrichment of proteins within substructures. For example, proteins may undergo low valency interactions with a large scaffold, such as segments of chromosomes or very long polymeric protein scaffolds [49]. These large scaffolds interact with proteins via spatially clustered binding sites (ICBS). Unlike in condensates, proteins in ICBS substructures do not phase separate. Full or partial FRAP assays yield similar recovery curves for substructures undergoing liquid-liquid phase separation or ICBS due to dynamic molecular interactions. To distinguish between condensates and ICBS substructures, half-FRAP (half-bleaching) method was developed [50]. This method is based on that proteins preferentially move and mix within the condensate and monitoring the maximum intensity decrease of the non-bleached half of the condensate (Fig 2C) [50]. In contrast, ICBS substructures do not show preferential movement of proteins inside or outside of substructures.

**Fig 2.**
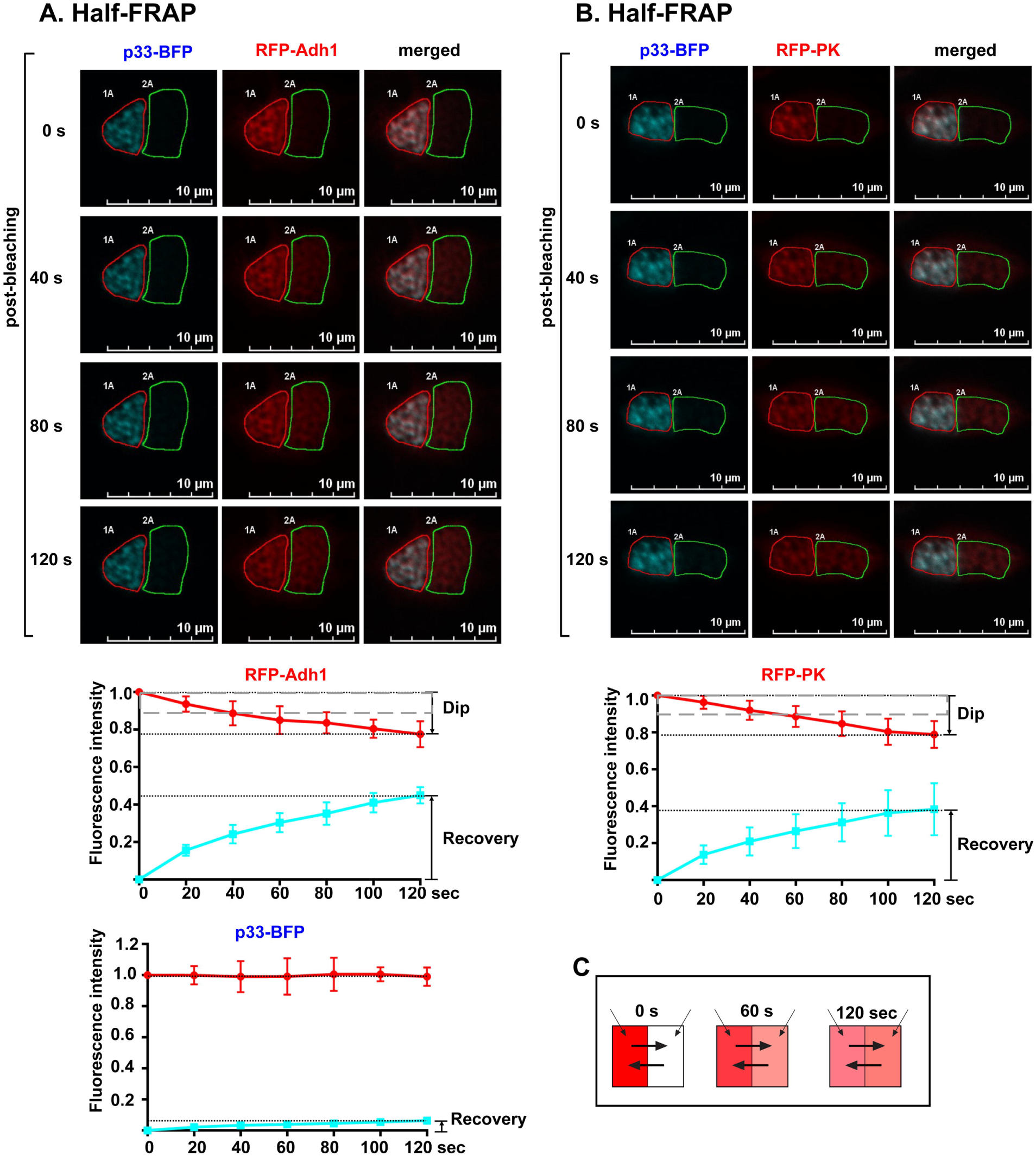
Half-FRAP experiments show condensate-like behavior of glycolytic proteins in VROs. (A) Half area of VROs containing RFP-Adh1 was photobleached in *N. benthamiana* cells. Then the fluorescent intensity was measured separately in the bleached (green circle) and the unbleached (red circle) halves of VROs for 120 sec. The extent of the fluorescent signal reduction in the unbleached half is shown as “Dip”. The fluorescent intensity of p33-BFP in the bleached and unbleached halves were quantified as control. (B) Comparable half-FRAP experiments with RFP-PK. (C) A scheme showing that the internal mixing of proteins within the condensate is predicted to be more efficient than with the surrounding cytosol. This protein mixing should increase the fluorescence signal in the bleached half while decrease the fluorescence signal in the unbleached half.

To test if the co-opted glycolytic proteins are present in condensate or ICBS substructures, we photobleached half of single VROs containing RFP-Adh1 (Fig 2A) or RFP-PK (Fig 2B), followed by measuring the intensity of fluorescent signals in the bleached versus unbleached halves of VROs. As expected, the fluorescent signal was partially recovered in the bleached half. However, the fluorescent signal was reduced by ∼23±6 % in the unbleached half of the VROs (Fig 2). This value is refered as “dip depth”. The half-FRAP data suggest that RFP-Adh1 and RFP-PK preferentially moved from the unbleached half to the bleached half. Previous studies showed that the dip depth value is >10% in condensates, whereas it is only 5-10% in ICBS substructures [50]. Thus, the data on the TBSV VROs indicate that the co-opted glycolytic enzymes are likely present in condensate substructures in VROs, and not in ICBS substructures.

To test if additional co-opted host proteins are also present in the vir-condensate portion of VROs, we selected Rpn11, which is proteasomal key protein interaction ‘hub’, a cytosolic protein [38, 51]. Rpn11 is a pro-viral host factor and it is efficiently co-opted into TBSV and CIRV VROs [38, 51, 52]. FRAP experiments revealed that Rpn11 was present in vir-condensates induced by TBSV p33 or CIRV p36 replication proteins (Fig 3A-B). We also confirmed that the co-opted Rpn11 was present in vir-condensates in TBSV-infected *N. benthamiana* cells (Fig 3C). Half-FRAP experiments showed that the RFP-Rpn11 signal was reduced by 24±5 % in the unbleached half of the VROs (Fig 3D). This high dip depth value also suggests that the co-opted Rpn11 is likely present in vir-condensate substructures, and not in ICBS substructures.

**Fig 3.**
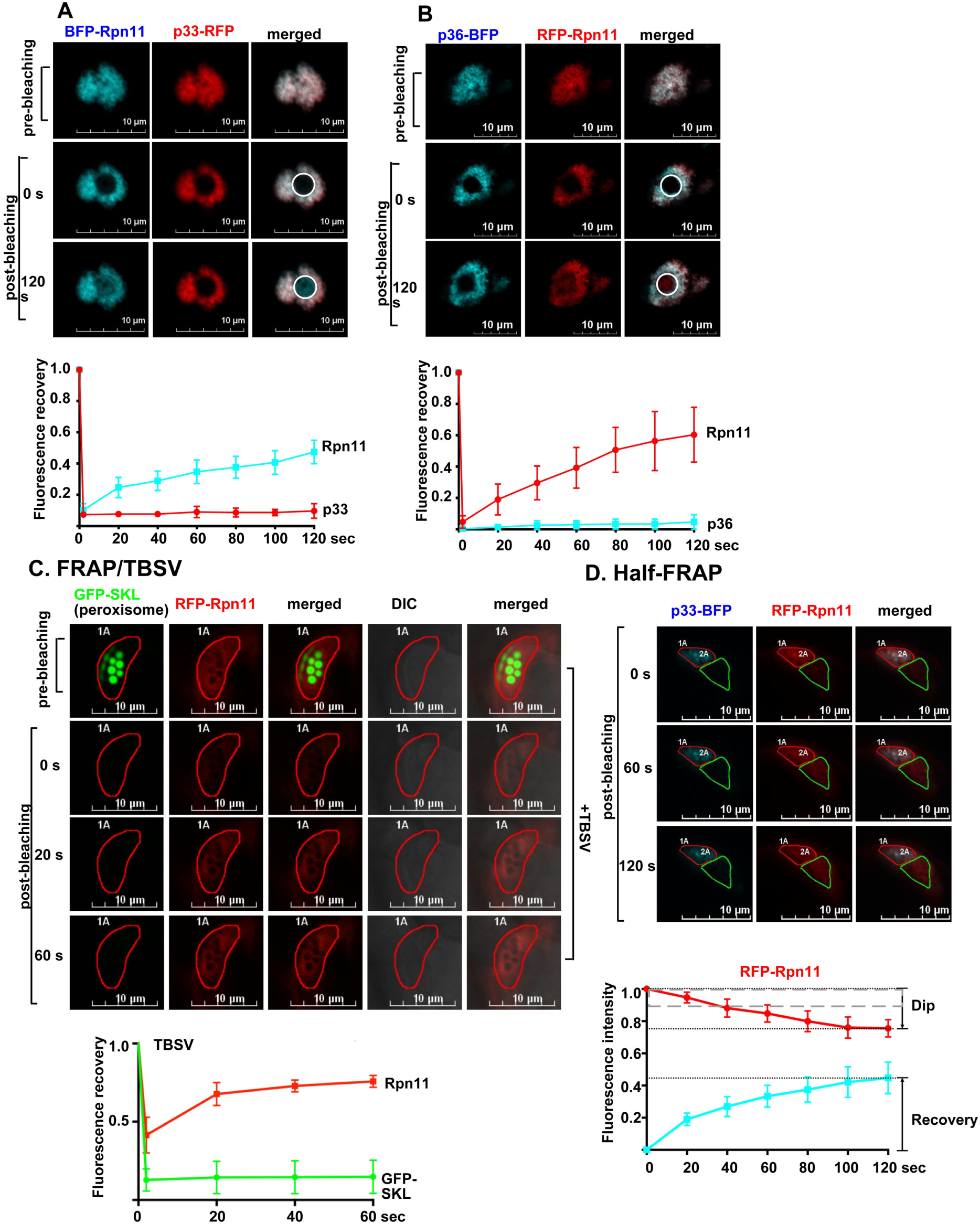
FRAP analysis of the co-opted non-glycolytic Rpn11 proteasomal protein in tombusvirus VROs. (A-C) FRAP analysis demonstrates the partial fluorescence recovery of either BFP-Rpn11 or RFP-Rpn11 after photobleaching within single VROs induced by either (A) TBSV p33-RFP, or (B) CIRV p36-BFP, or (C) TBSV infection in *N. benthamiana*. The clustered peroxisomes in a single VRO were marked with GFP-SKL peroxisomal marker. Scale bars represent 10 μm. The graphs at the bottom panel show the extent of fluorescence recovery of Rpn11 within VROs at a given time point. See further details in Fig 1. (D) Half-FRAP analysis of VROs containing RFP-Rpn11 in *N. benthamiana* cells. The fluorescent intensity was measured separately in the bleached and the unbleached halves of VROs for 120 sec. See further details in Fig 2.

The FRAP recoveries for the co-opted host proteins are generally slow and incomplete, indicative of gel-like condensates. Highly dynamic condensates (FUS, TDP43, etc.) can recover in seconds [41, 54], while the vir-condensates identified in this study are less dynamic. To test if small proteins could move faster within the vir-condensates, we chose the 10 kB SUMO1 protein, which is enriched in VROs (Fig S4). FRAP assay showed much faster and close to full recovery of the fluorescence signal for RFP-SUMO1 in VROs (Fig S4), suggesting that smaller molecules could move more rapidly than the larger proteins within the vir-condensates.

The distributions of the co-opted glycolytic PK and the fermentation Pdc1 and Adh1 enzymes and Rpn11 protein within VROs were partially affected by treatment with 10% 1,6-hexanediol (Fig S5A-D), which dissolves condensates through disrupting weak hydrophobic interactions between proteins [39, 45]. This indicates that electrostatic and hydrophobic interactions between proteins contribute to vir-condensates formation. Comparable 1,6-hexanediol treatment led to disapprerance of P bodies in *N. benthamiana* cells, marked with GFP-Dcp1 (Fig S5E), which is known to form liquid-like condensates in plant cells [53].

### TBSV p33 and CIRV p36 replication proteins form droplets *in vitro*

To demonstrate if TBSV p33 and the CIRV p36 replication proteins could induce phase separation, we performed *in vitro* droplet formation experiments as shown in Fig 4. These experiments demonstrated that the purified TBSV eGFP-p33C and CIRV eGFP-p36C replication proteins formed round droplets under physiological conditions in the presence of 2.5% or more PEG8000 crowding agent after 10 min incubation at room temperature (Fig 4). The sizes and numbers of droplets formed were dependent on protein and PEG8000 concentrations. FRAP experiments showed that the fluorescence signal in the droplet formed by TBSV eGFP-p33C did not recover (Fig 5B). The droplets were stable and did not fuse with each other (Fig 5C). These data indicate that droplets formed by the C-terminal fragment, which is exposed to the cytosol [55, 56], of TBSV p33 replication protein, are in gel-like, not liquid-like forms.

**Fig 4.**
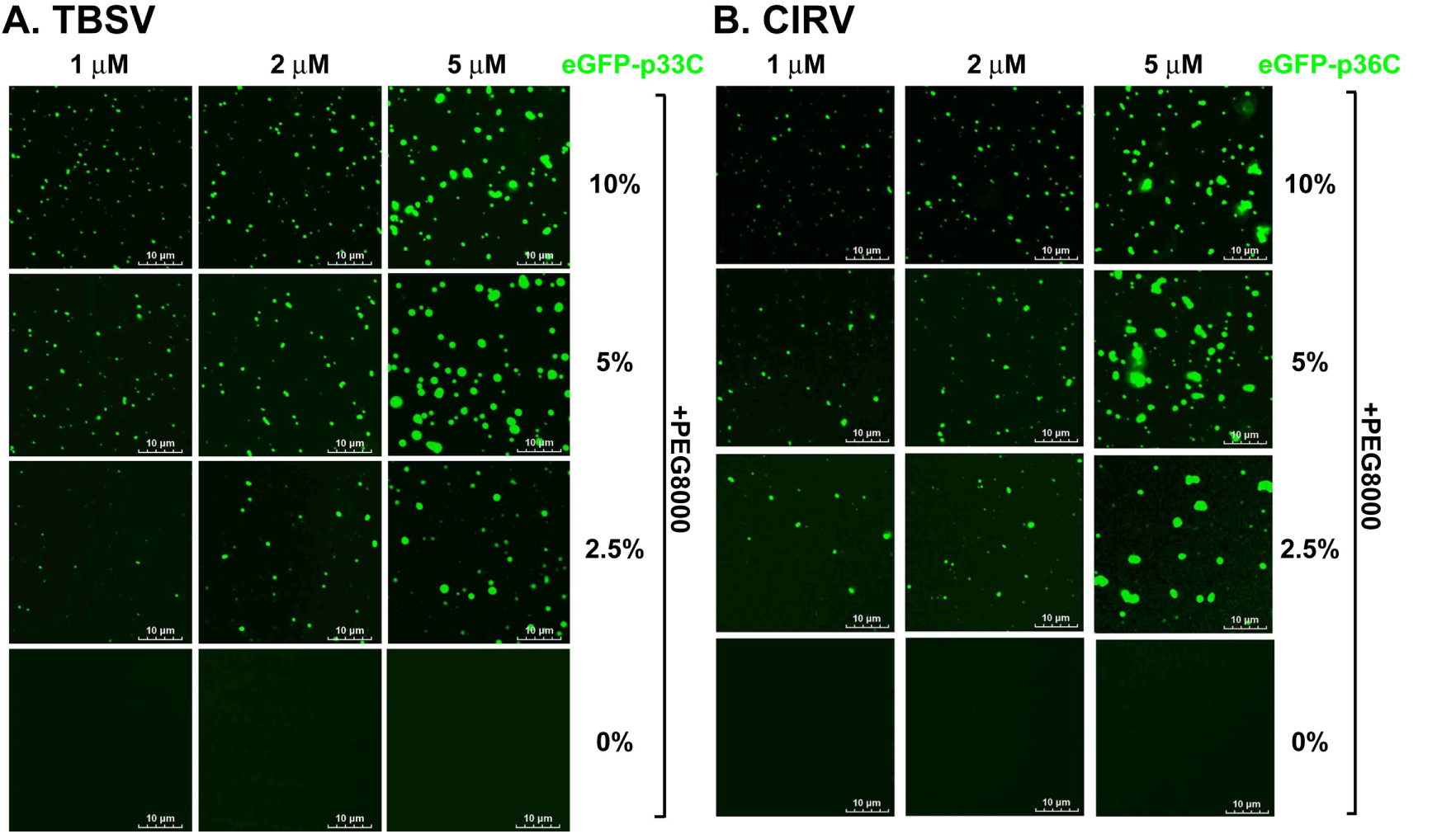
Droplet formation by the tombusvirus replication proteins *in vitro*. (A) The purified recombinant cytosol-exposed C-terminal half of p33 (eGFP-p33C) was incubated in the presence of increasing amounts of PEG8000. Confocal imaging shows the droplet formation. Scale bars represent 10 μm. (B) Droplet formation by the purified recombinant CIRV eGFP-p36C. See further details in panel A.

**Fig 5.**
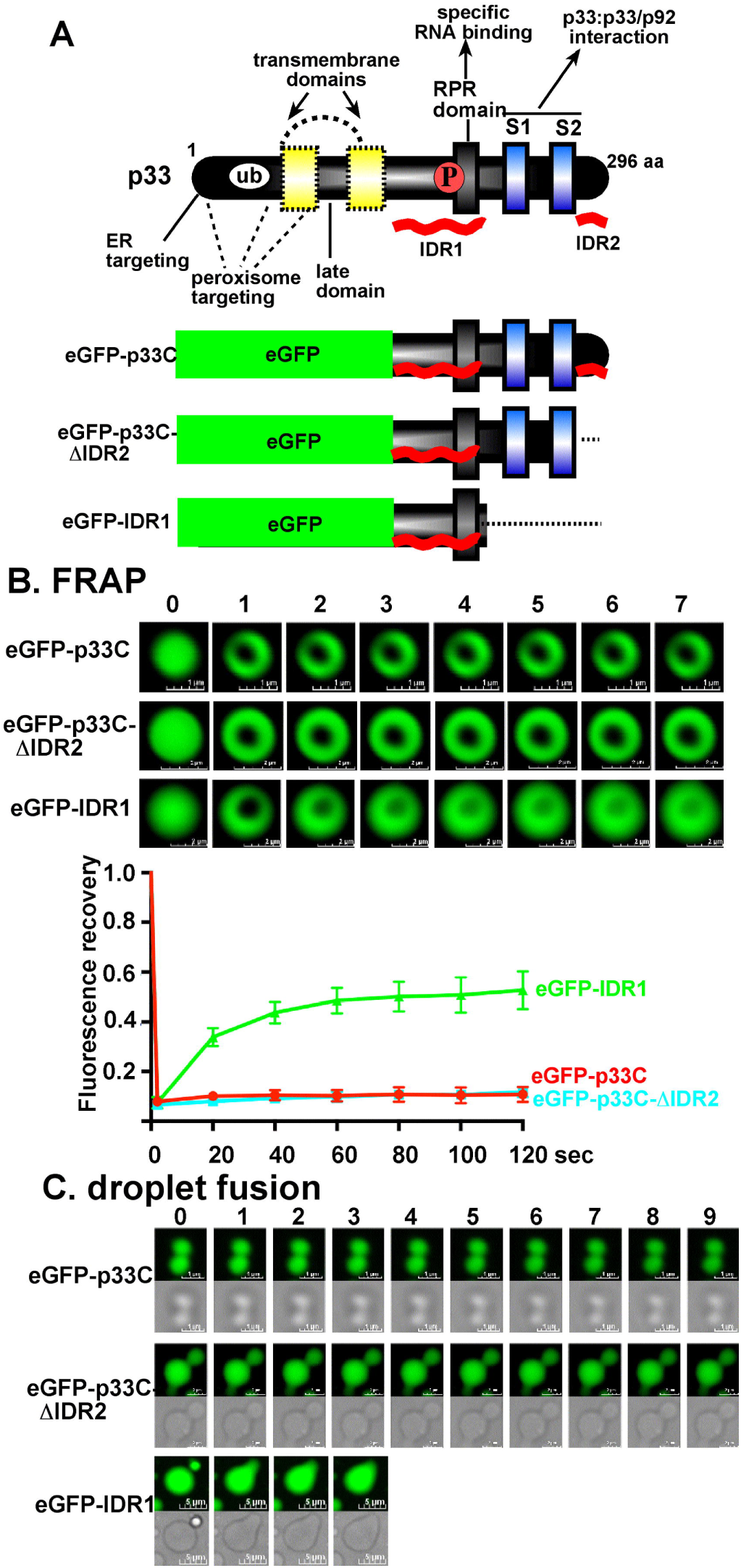
Droplets formed by the TBSV p33 replication protein *in vitro* show gel-like properties. (A) Scheme of the TBSV p33 replication protein with the known domains. The curved red lines indicate the two IDR regions in p33. The two deletion mutants are also shown. (B) FRAP analysis shows the lack of fluorescence recovery of either eGFP-p33C or eGFP-p33C-ΔIDR2 signals after photobleaching within single droplets. The bottom panel shows the fluorescence recovery of eGFP-IDR1 signal. The images were taken sequentially at every 20 sec after photobleaching. FRAP analysis was conducted on 8 independent droplets per expreiment. The graph shows the extent of fluorescence recovery. (C) Confocal and DIC images of droplet fusion experiments show the lack of fusion between droplets formed by either eGFP-p33C or eGFP-p33C-ΔIDR2. The bottom panel shows that the droplets formed by eGFP-IDR1 fused rapidly. The images were taken sequentially at every 6.5 sec. Each experiment was repeated three times.

Treatment with 10% 1,6-hexanediol, which dissolves condensates through disrupting weak hydrophobic interactions between proteins [39, 45] showed that the droplet formed by TBSV p33C were partially sensitive in the presence of 2.5% PEG8000, whereas TBSV p33C droplets were insensitive in the presence of 10% PEG8000 (Fig S6). These data suggest that the protein interaction between p33C molecules is likely based on both electrostatic and hydrophobic interactions.

To identify what region in TBSV p33 protein or in CIRV p36 organizes phase separation/droplet formation, we deleted the very C-terminal intrinsically disordered region (IDR2, Fig S7A-B) and the p33:p33/p92 interaction domain from eGFP-p33C (eGFP-IDR1, Fig 5A). The latter construct contained only the long IDR1 region, including the RPR domain, which is a specific RNA binding region [23, 57]. Deletion of IDR2 (eGFP-p33C-ΔIDR2) resulted in larger droplets than the droplets formed by eGFP-p33C (Fig S7C). FRAP-based experiments showed that the fluorescence signals in droplets formed by eGFP-p33C-ΔIDR2 did not recover, similar to eGFP-p33C (Fig 5B). These data suggest that the IDR2 region has inhibitory effect on droplet expansion. We found that the TBSV eGFP-IDR1 protein formed droplets *in vitro*, suggesting that the IDR1 region within p33 replication protein is critical for phase separation/droplet formation. The number of droplets formed by eGFP-IDR1 was much less than those of eGFP-p33C, but grew bigger after 30 min incubation (Fig S7C). Moreover, FRAP-based experiments showed that the fluorescence signals in TBSV eGFP-IDR1 droplets recovered rapidly (Fig 5B). In addition, the eGFP-IDR1-induced droplets fused rapidly with one another (Fig 5C). These results suggest that the IDR1 region of p33 induces liquid-like droplets, unlike the more solid, gel-like droplets induced by p33C. Thus, the p33:p33/p92 interaction domain in p33 replication protein likely contributes to droplet formation by strengthening the protein-protein interactions, which seem to be weak between the IDR1-containing proteins. Deletion of portion of the IDR1 sequence in p33C led to protein aggregation *in vitro* (Fig S7D), suggesting that the IDR1 region is critical for droplet formation. The CIRV p36-based experiments also showed that deletion of the IDR1 region in eGFP-p36C-ΔIDR1 blocked droplet formation, whereas the CIRV IDR1 region induced droplets (eGFP-IDR1, Fig S7A). Altogether, the *in vitro* droplet formation experiments demonstrated that both TBSV p33 and CIRV p36 use the IDR1 region for inducing phase separation and the C-terminal the p33:p33/p92 interaction domain further facilitates phase separation and likely contribute to stabilization of the vir-condensates.

### Glycolytic and fermentation enzymes partition to droplets formed by TBSV p33 or CIRV p36 replication proteins *in vitro*

The ability of TBSV p33 and CIRV p36 replication proteins to form droplets *in vitro* raises the possibility that these replication proteins could promote partitioning of the co-opted host proteins to p33/p36 droplets. This was tested using an *in vitro* co-droplet formation assay. Purified TBSV eGFP-p33C (5 μM) or CIRV eGFP-p36C (5 μM) were co-incubated with purified RFP-Adh1 (0.5 and 5 μM) in the incubation buffer (physiological salt concentration and pH) containing 2.5% PEG8000 on a glass bottom dish at room temperature for 30 min. Analysis with a confocal microscope confirmed that RFP-Adh1 was partitioned to p33/p36 droplets (Fig 6). As a control, eGFP-p33C or CIRV eGFP-p36C did not promote the partitioning of purified RFP to droplets (5 μM) under the same conditions (Fig 6). RFP-Adh1 did not form droplet (5 or 10 μM) or co-droplet with eGFP (5 μM) (Fig 6A) under the same conditions, thus demonstrating that the TBSV p33 or CIRV p36 replication proteins, not Adh1, organize the phase separation *in vitro*. Similar experiments with plant NbPgk1 (5 or 0.5 μM), and PK or yeast ScPgk1, ScPdc1 (5 or 0.5 μM), ScAdh1, and Cdc19 (the yeast PK homolog) (Fig S8) and Rpn11 or ScRpn11 (Fig S9) also demonstrated that p33C or p36C promoted the partitioning of the above proteins to droplets. Droplet formation assay demonstrated that high protein concentration and 10% PEG8000 promoted droplet formation by Adh1, Pgk1, PK and Rpn11 *in vitro* (Fig S10), suggesting that these cytosolic proteins can undergo liquid-liquid phase separation under suitable conditions. Altogether, TBSV p33 and CIRV p36 replication proteins can undergo phase separation and the co-opted cytosolic host factors partition to p33C/p36C droplets *in vitro*.

**Fig 6.**
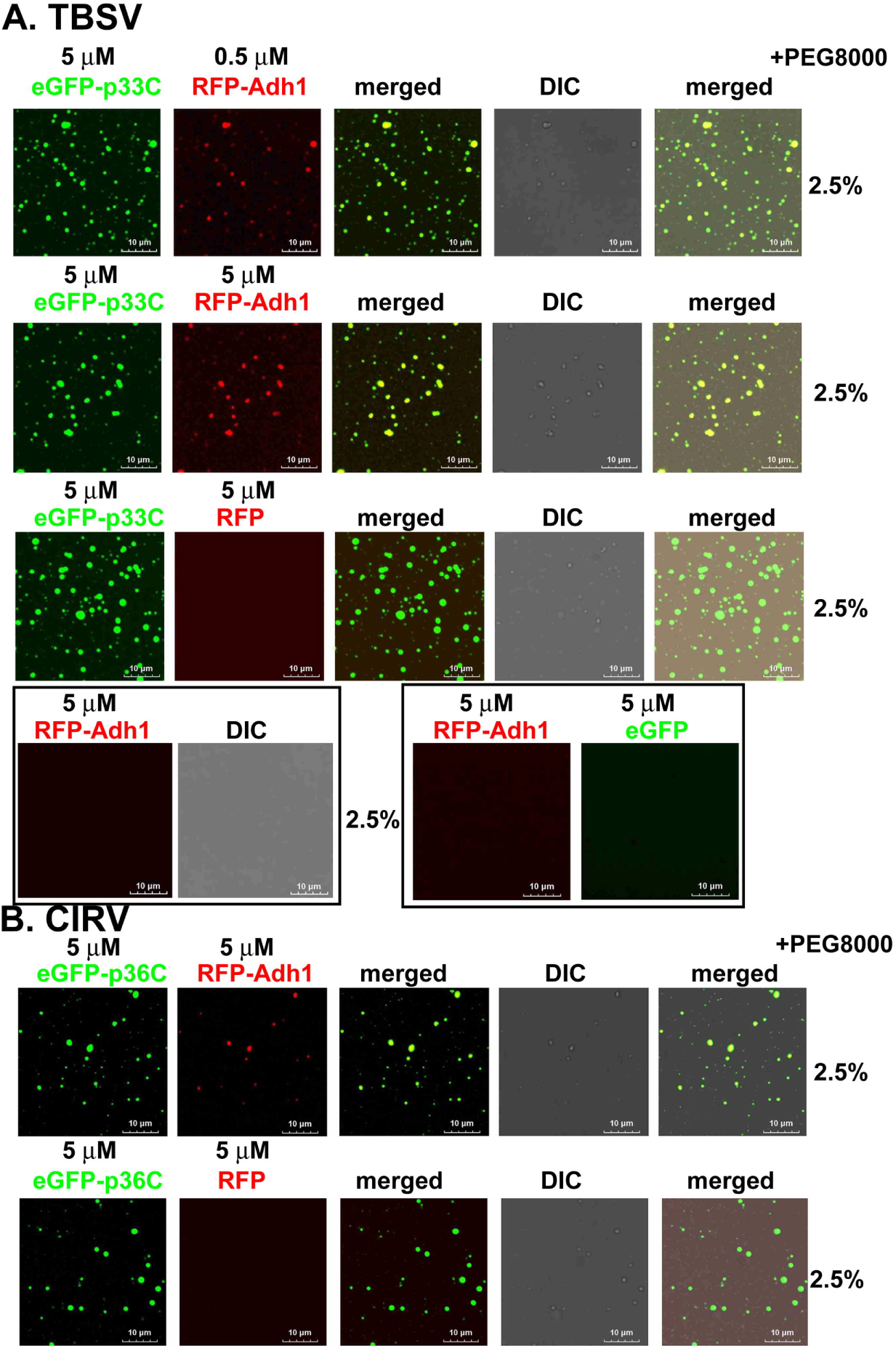
The tombusvirus replication proteins induce the partitioning of purified Adh1 fermentation enzyme to droplets *in vitro*. (A) Confocal images show the partitioning of the purified RFP-Adh1 to droplets formed by purified recombinant eGFP-p33C in the presence of 2.5% PEG8000. Note that RFP did not partition to GFP-p33C droplets and no droplet was observed between RFP-Adh1 and eGFP under the same conditions. (B) Confocal images show the partitioning of the purified RFP-Adh1 to droplets formed by purified recombinant CIRV eGFP-p36C in the presence of 2.5% PEG8000. Note that the purified RFP did not partition to CIRV eGFP-p36C droplets in the presence of 2.5% PEG8000. Scale bars represent 10 μm. Each experiment was repeated three times.

### The IDR1 region in TBSV p33 and CIRV p36 replication proteins is critical for VRO formation and function

To test if the IDR1 region, which drives phase separation *in vitro* (Fig 5), is required for VRO formation, we expressed TBSV eGFP-p33 lacking the IDR1 region (eGFP-p33-ΔIDR1) or CIRV eGFP-p36-ΔIDR1 in *N. benthamiana*. Confocal microscopy revealed that eGFP-p33-ΔIDR1 or eGFP-p36-ΔIDR1 formed small puncta, unlike the large VRO formed by either WT eGFP-p33 or eGFP-p36 (Fig 7). As expected, eGFP-p33-ΔIDR1, which carries the peroxisomal targeting signals [55, 58], was co-localized with the peroxisomal marker protein in *N. benthamiana* (Fig 7A). Both CIRV eGFP-p36-ΔIDR1 and the WT eGFP-p36 were co-localized with the mitochondrial marker in *N. benthamiana* (Fig 7B). Confocal imaging showed that RFP-PK or RFP-Adh1 were infrequently enriched in the eGFP-p33-ΔIDR1 puncta (Fig 8B-C). Deletion of IDR1 in CIRV p36 replication protein (eGFP-p36-ΔIDR1) led to formation of small puncta, which was co-localized with RFP-PK in a few eGFP-p36-ΔIDR1 puncta, but not all (Fig 8D). Expression of a C-terminally truncated p33 (eGFP-p33-sIDR1) containing the N-terminal sequences and a portion of IDR1 or the entire IDR1 (eGFP-p33-IDR1) resulted in VRO-like structure formation (Fig 8A-B). RFP-PK was enriched in the eGFP-p33-sIDR1 or eGFP-p33-IDR1-induced VRO-like structures (Fig 8B). FRAP experiments revealed that p33-sIDR1-BFP or p33-IDR1-BFP -induced VROs contain vir-condensate because of the partial recovery of fluorescence signal of RFP-PK (Fig 8E). The fluorescence signal of RFP-PK recovered faster and to a higher level in p33-sIDR1-BFP or p33-IDR1-BFP -induced VROs than in the WT p33-induced VROs (Fig 8E), indicating more liquid-like behavior of the vir-condensates induced by p33-sIDR1 or p33-IDR1.

**Fig 7.**
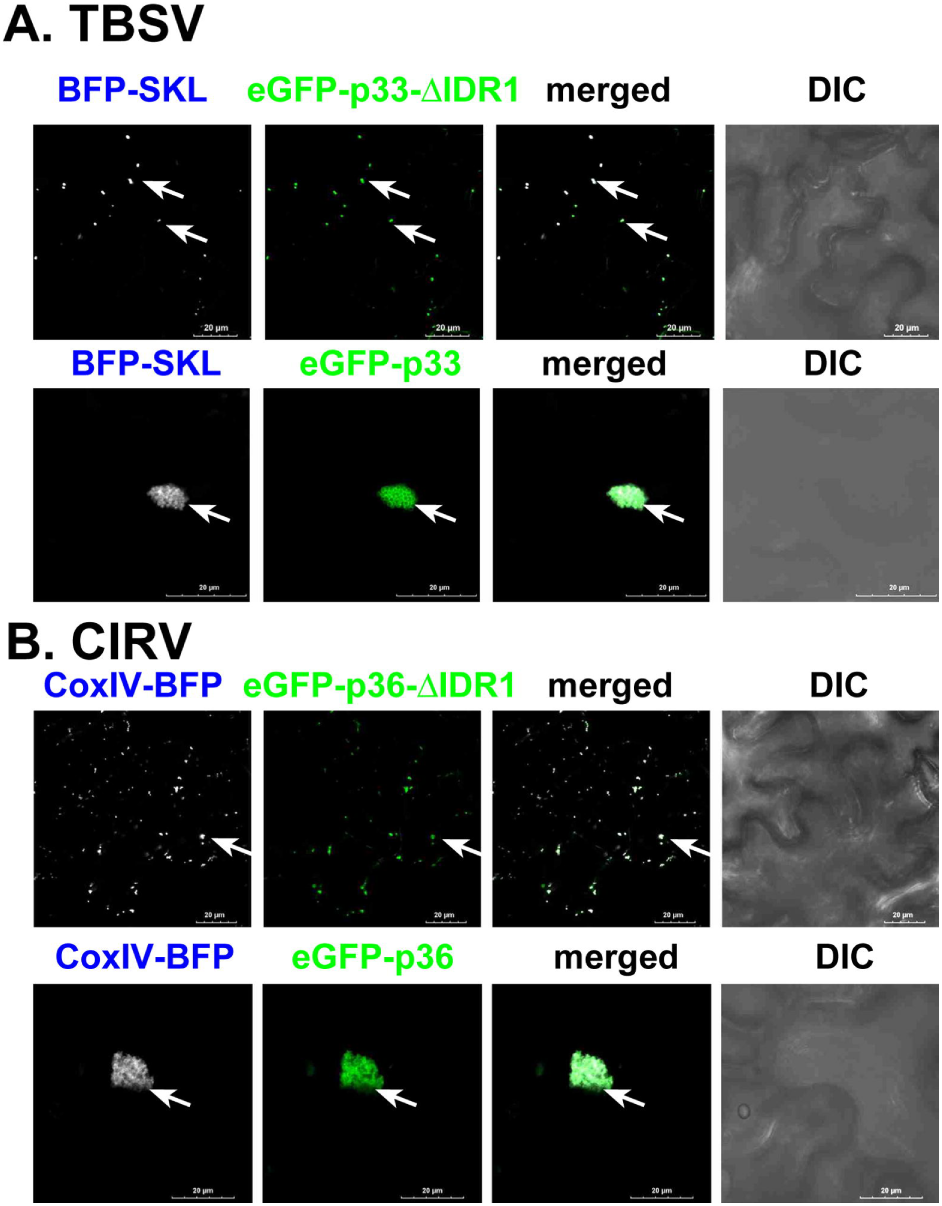
The IDR1 region in the tombusvirus replication proteins is required for VRO formation in plants. (A) Confocal images show the lack of peroxisome clustering, which is a hallmark of TBSV VRO formation, in *N. benthamiana* cells expressing TBSV p33 lacking IDR1 (eGFP-p33-ΔIDR1). The control wt eGFP-p33 expression induces VRO formation. BFP-SKL marks the peroxisomes. (B) Confocal images show the lack of mitochondria clustering, which is a hallmark of CIRV VRO formation, in *N. benthamiana* cells expressing CIRV p36 lacking IDR1 (eGFP-p36-ΔIDR1). CoxIV-BFP is the mitochondrial marker protein. Scale bars represent 20 μm. Each experiment was repeated three times.

**Fig 8.**
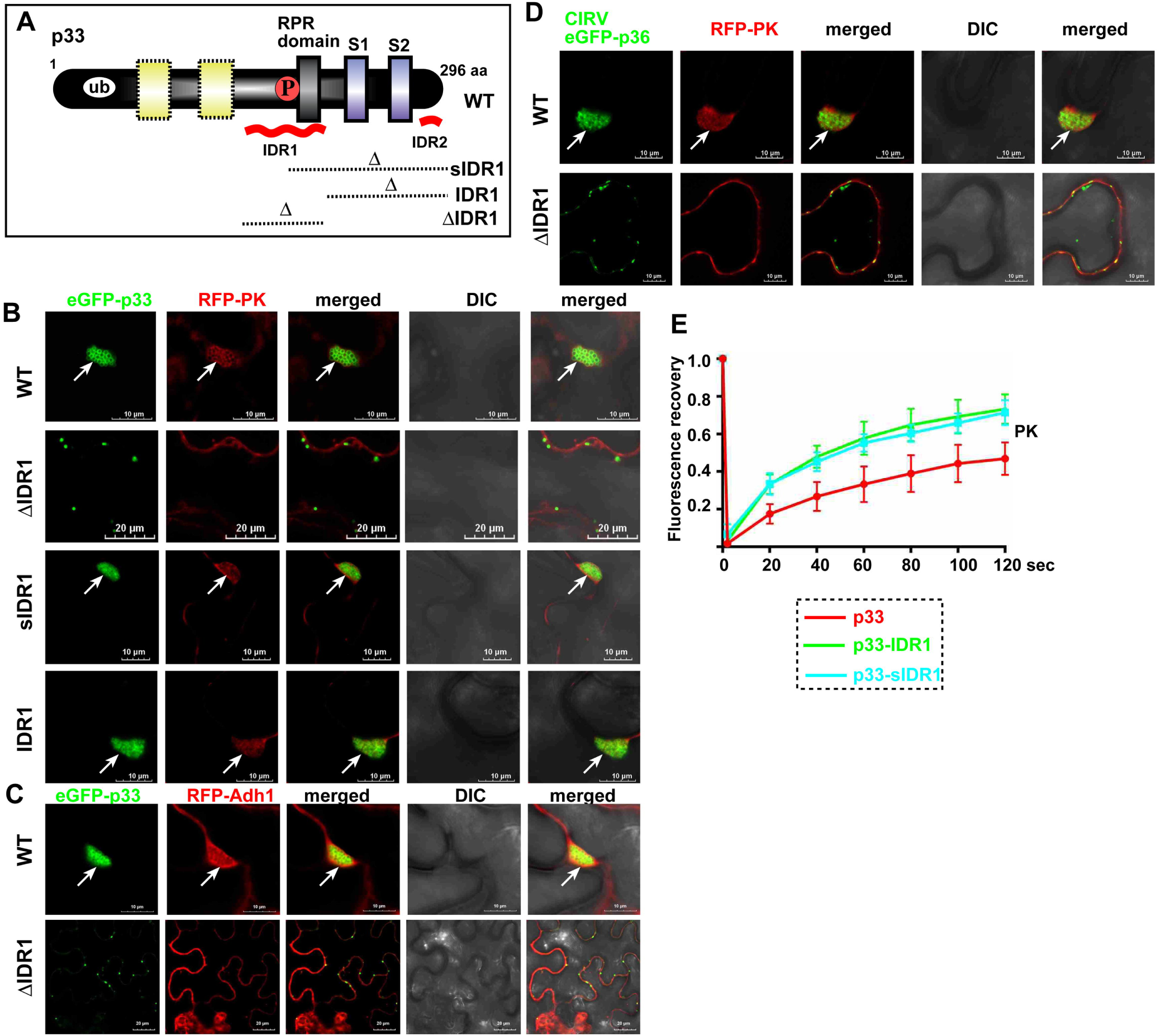
The IDR1 region in the tombusvirus replication proteins is needed for enrichment of glycolytic and fermentation enzymes in VROs. (A) A scheme of the p33 deletion mutants used. See Fig 5A for further details. (B) Colocalization of pyruvate kinase (RFP-PK) with eGFP-p33 or mutants in *N. benthamiana* cells. Arrows point at individual VROs. Scale bars represent 10 or 20 μm as shown. (C) Colocalization of RFP-Adh1 with eGFP-p33 or mutant (eGFP-p33-ΔIDR1) in *N. benthamiana* cells. See further details in panel B. (D) Colocalization of RFP-PK with CIRV eGFP-p36 or mutant (eGFP-p36-ΔIDR1) in *N. benthamiana* cells. See further details in panel B. Each experiment was repeated three times. (E) FRAP analysis of fluorescence recovery of RFP-PK in p33-BFP or mutant-induced VROs in *N. benthamiana* cells. 6-to-8 VROs from three independent plants were tested in each experiment.

Charged amino acids are frequently involved in phase separation [54, 59] and the IDR1 region in p33 contains a highly charged segment (Fig 9A). We mutated this segment to eliminate the 7 charged amino acids, creating eGFP-p33-IDR1m1 (Fig 9A). The purified eGFP-IDR1m1 produced 3-fold less and much smaller droplets than eGFP-IDR1 (Fig 9B). Confocal microscopy studies revealed that eGFP-p33-IDR1m1 induced much smaller VROs than WT eGFP-p33-IDR1 in *N. benthamiana* cells (Fig 9C). These data indicate that the charged segment of p33 IDR1 contributes to vir-condensate formation.

**Fig 9.**
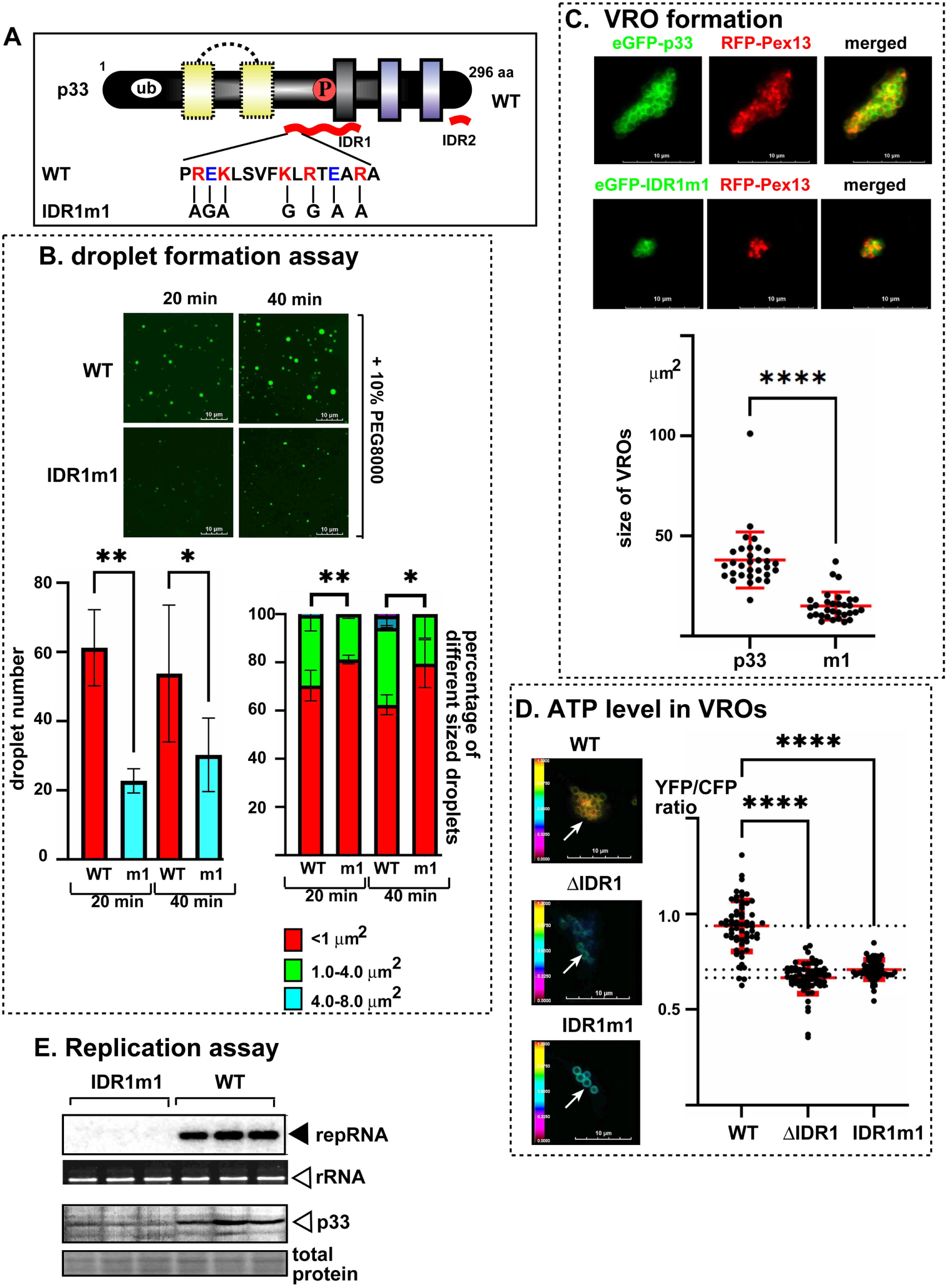
The effects of IDR1 mutations on the functions of p33 replication protein. (A) A scheme of the p33 IDR1 mutant (p33-IDR1m1). (B) Top images: confocal images of droplets formed by either WT eGFP-IDR1 or eGFP-IDR1m1. Note that we used the same eGFP-IDR1 as shown in Fig 5A. Bottom left panel shows the reduced droplet number whereas bottom right panel shows the distribution of doplet size supported by eGFP-IDR1m1. Red box indicates droplet size of <1.0 μm^2^, green box represents 1.0-4.0 μm^2^, blue box represents the % of droplets of 4.0-8.0 μm^2^. (C) Confocal images show the reduced size of VROs formed in *N. benthamiana* cells expressing eGFP-p33-IDR1m1 versus WT eGFP-p33. Scale bars represent 10 μm. The sizes of VROs are measured in μm^2^. (D) Reduced ATP production within the VROs formed by p33-IDR1m1 or p33-ΔIDR1 in *N. benthamiana* cells, which expressed the p33 or mutants fused with an ATP biosensor (ATeam^YEMK^). The more intense FRET signals are white and red (between 0.5 to 1.0 ratio), whereas the low FRET signals (0.1 and below) are light blue and dark blue. We show the quantitative FRET values (obtained with ImageJ) for numbers of samples in the graph. (E) Top image: Northern blot analysis demonstrates decreased TBSV replicon (+)RNA accumulation in yeast. The His_6_-tagged WT p33 or p33-IDR1m1, p92^pol^ and DI-72 replicon RNA were co-expressed from plasmids. The accumulation level of replicon RNA was normalized based on 18S rRNA levels (second panel). Bottom panels: The accumulation of His_6_-p33 or His_6_-p33-IDR1m1 is measured by western blotting and anti-His_6_ antibodies. Each experiment was repeated three times.

The major pro-viral roles of the co-opted glycolytic and fermentation enzymes are to produce ATP locally within the VROs to support the large energy need of rapid virus replication [34, 35, 37]. To test if vir-condensate formation is needed for ATP production within the VROs, we used an ATP-biosensor module fused to TBSV p33 or mutants, which targets the fusion protein to VROs. Briefly, p33-ATeam measures ATP level within the VROs based on conformational change in the ε subunit of the bacterial F_0_F_1_-ATP synthase after ATP binding [37, 60]. The ε subunit in ATP-free form is in extended conformation (low FRET value), whereas the ATP bound conformation is placing the CFP and YFP in proximal position in p33-ATeam, leading to enhanced FRET signals (Fig S11).

We found that VROs induced by either p33-ΔIDR1 or p33-IDR1m1 with defective vir-condensates contained significantly less ATP than VROs induced by the full-length p33 (Fig 9D), suggesting that vir-condensate formation is critical for local ATP production within the VROs. The sizes of the VROs formed by WT p33 did not affect ATP generation within the VROs (Fig S12). Moreover, p33-IDR1m1 did not support replication of TBSV replicon RNA in yeast cells (Fig 9E). Altogether, the obtained data support the model that the IDR1 region in TBSV p33 or CIRV p36 replication proteins plays a critical role in organizing vir-condensates, ATP production and in VRO formation and functions.

### The co-opted ER membranes and subverted actin filaments form meshworks in and around the vir-condensates

A fraction of ER membranes is co-opted by tombusviruses via the formation of membrane contact sites (vMCS) between the ER and the peroxisomes (in case of TBSV) or mitochondria (CIRV) [27, 29, 30]. To test if the ER membranes are associated with the vir-condensates, we co-expressed ER marker proteins (either GFP-HDEL or BFP-CTT) [61] with RFP-PK and p33-BFP or BFP-Adh1 and p33-RFP in *N. benthamiana* cells. Interestingly, confocal images showed that the ER formed a meshwork in and around the vir-condensates and VROs (Fig 10A). The ER membranes were highly enriched around and within VROs, whereas ER enrichment was not detectable in the vicinity of peroxisomes in the absence of TBSV p33 (Fig S13B). Similar enrichment of ER meshwork was observed in and around the vir-condensates and VROs induced by the CIRV p36 replication protein (Fig 10A). We confirmed with additional co-opted glycolytic proteins and Rpn11 that the ER meshwork forms around the vir-condensates and VROs (Fig S14).

**Fig 10.**
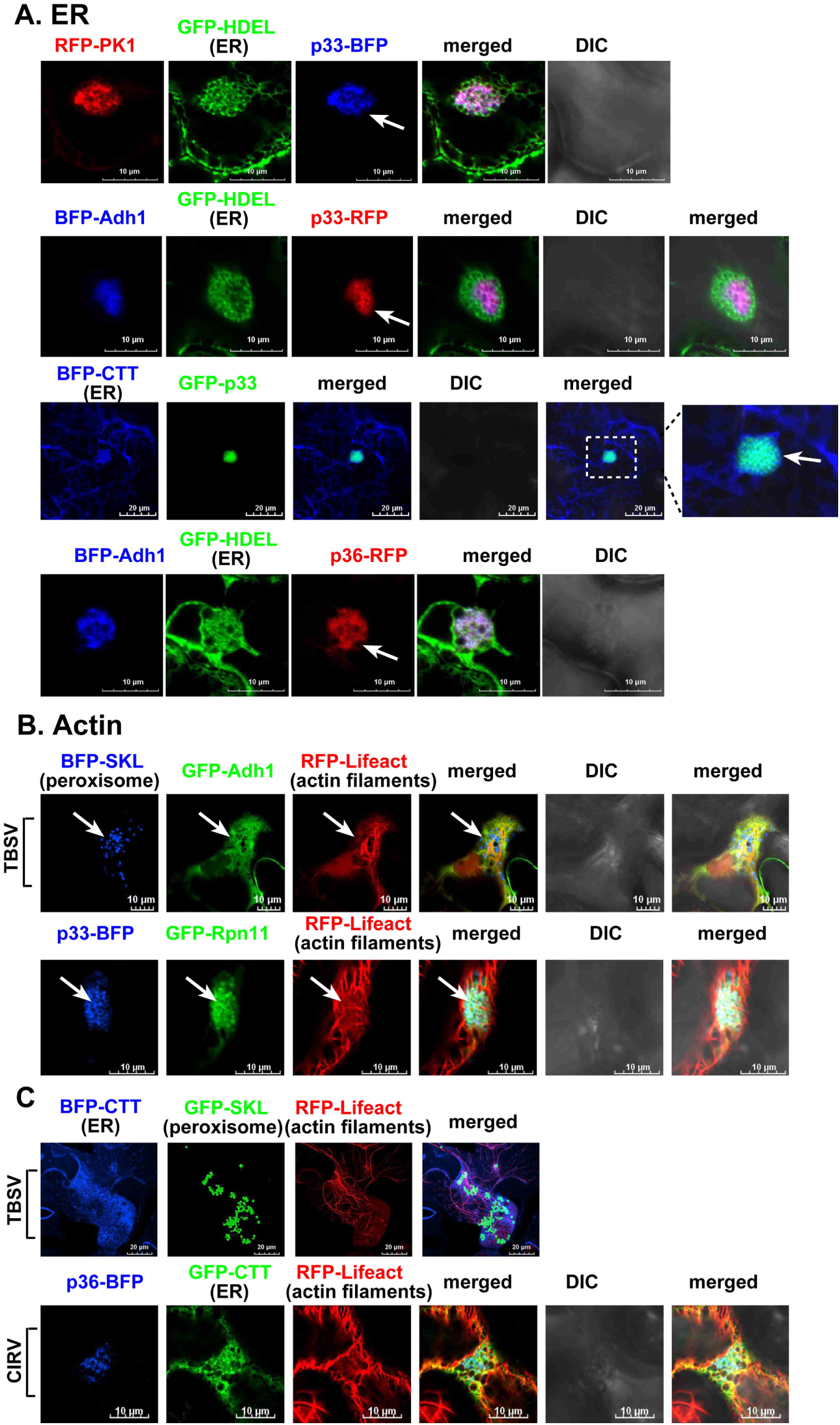
The co-opted actin filaments and the ER membranes form meshworks around the VROs. (A) Transient expression of GFP-HDEL or BFP-CTT was used to visualize the ER meshwork formed around VROs marked by TBSV p33 or CIRV p36 in *N. benthamiana* cells infected with either TBSV or CIRV. Arrows point at individual VROs. (B) Transient expression of RFP-Lifeact was used to visualize the co-opted actin filament meshwork formed around VROs marked by TBSV p33 or CIRV p36 in *N. benthamiana* cells infected with either TBSV or CIRV. Arrows point at individual VROs. (C) Visualization of the ER and actin meshwork around VROs marked by GFP-SKL peroxisomal marker or CIRV p36-BFP in *N. benthamiana* cells infected with either TBSV or CIRV. Scale bars represent 10 μm. Each experiment was repeated three times.

The actin network is co-opted for tombusvirus replication, which facilitates the VRO assembly [38, 62, 63]. To examine if the actin filaments are associated with the vir-condensates, we expressed the actin filament marker (RFP-Lifeact), p33-BFP and either GFP-Adh1 or GFP-Rpn11 in *N. benthamiana* cells infected with TBSV. Confocal images showed that the actin filaments were present in the vir-condensates and the VROs (Fig 10B). The co-opted ER membranes and actin filaments formed an entagled meshwork in and around the VROs induced by TBSV or CIRV infections in *N. benthamiana* cells (Fig 10C and Fig S15).

## Discussion

Similar to many other plant and animal viruses [1, 3, 10, 24, 64–66], tombusvirus VROs contain a large number of tiny spherules (small invaginations) in co-opted membranes, i.e. peroxisomal membranes for TBSV and mitochondrial membranes for CIRV. These semi-closed spherules harbor replication complexes, including the two viral replication proteins and the viral RNA, and support viral RNA replication [16, 18]. However, tombusviruses co-opt numerous host proteins, many of them cytosolic, not membrane-bound, such as the glycolytic enzymes. Here, we showed evidence that the co-opted glycolytic and fermentation enzymes, plus the proteasomal Rpn11 pro-viral host factors were sequestered into a vir-condensate substructure, which constituted an integral part of the membranous VROs [24]. FRAP-based experiments revealed that the fluorescence signals of the membrane-bound TBSV p33 and CIRV p36 did not recover after photobleaching. On the other hand, fluorescence signals of the co-opted glycolytic and fermentation enzymes, plus Rpn11 cytosolic protein, were partially recovered within the VROs. The relatively slow and partial recovery of fluorescence signals after photobleaching indicates that the vir-condensates seem to be in a gel-like form. These findings suggest that the tombusvirus VROs contain two distinct substructures: one consists of the subverted cellular membranes and the other is the vir-condensate, which allows relatively slow movement of co-opted host protein within the vir-condensate. It is important to note that p33/p36 replication proteins induce and connect the two substructures, which co-exist within the VRO (Fig 11) and both are critical for VRO functions.

**Fig 11.**
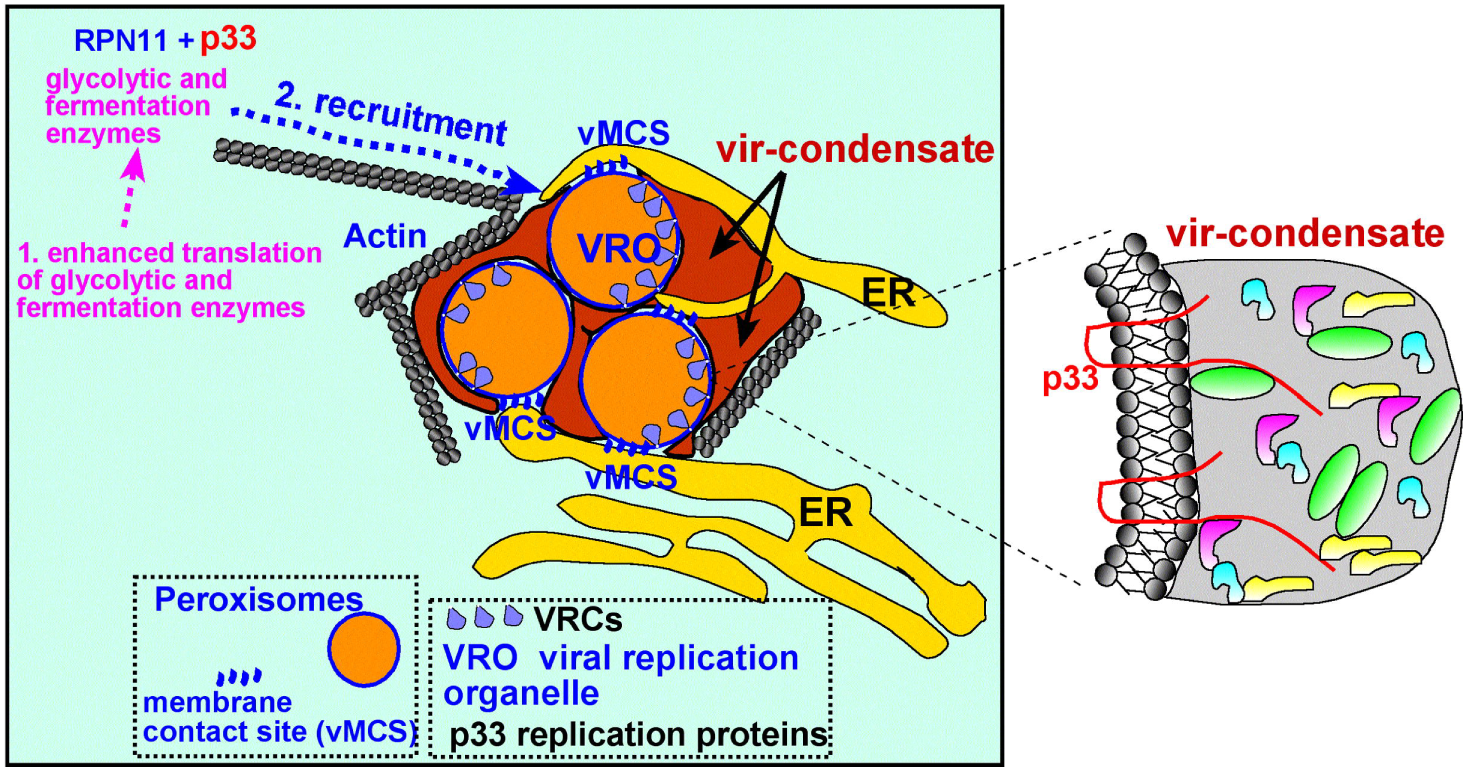
A model on the role of phase separation in TBSV VRO formation. We propose that TBSV p33 replication protein induces liquid-liquid phase separation of the co-opted, highly concentrated glycolytic and fermentation enzymes and Rpn11, forming unique gel-like vir-condensates associated with VROs, which consist of clustered peroxisomes. The enlarged panel shows that TBSV p33 replication protein is present both in the membranous part of VRO via the N-terminal two transmembrane domains and in the vir-condensate via the C-terminal (cytosol-exposed) region. We propose that co-opted ER membranes and actin filaments provide physical barriers, which facilitate in keeping the virus-induced vir-condensate (in red) and the co-opted clustered peroxisomes within the VRO boundary. A major function of the vir-condensate containing the co-opted glycolytic and fermentation enzymes is to efficiently produce ATP *in situ* within the VROs. The p33 replication protein is the main organizer by connecting the co-opted membranous substructure and vir-condensate within VRO. CIRV p36 replication protein has similar functions to the TBSV p33 replication protein, except the VROs consist of clustered mitochondria. Abbreviations: VRC: viral replicase complex; vMCS: virus-induced membrane contact site.

The vir-condensates were also formed during full tombusvirus infections in *N. benthamiana* cells. The formation of special vir-condensate facilitated enrichment and sequestration of co-opted cytosolic enzymes, which were associated with the membranous VROs during the entire replication process [34–37, 48]. We propose that the vir-condensate is likely associated with and surround the “core” clustered peroxisomes or mitochondria (Fig 11). Sequestration of cytosolic enzymes by vir-condensates likely occurs in direct competition with cellular condensates that sequester these same enzymes (e.g., GAPDH). A recent study found that the antiviral CASC3 condensates efficiently sequester tombusvirus host factors like GAPDH, Vps29, and HSP70 [67].

FRAP assay showed that the co-opted glycolytic and fermentation enzymes and Rpn11 [35–38, 48, 51] were present in vir-condensates in VROs. TBSV p33 and CIRV p36 replication proteins effectively sequestered/concentrated these host proteins in a confined space. We propose that when the amounts of these co-opted proteins reach the threshold of phase separation, then p33/p36 replication proteins organize liquid-liquid phase separation of co-opted highly concentrated host proteins within VROs through multivalent interactions (Fig. 11). Since the multivalent interactions between these co-opted cytosolic proteins and p33/p36 could be dynamic and reversible, these cytosolic proteins can move within the vir-condensate. However, p33/p36 are membrane proteins anchored to peroxisomes or mitochondria and their mobility is much less dynamic than cytosolic proteins within the vir-condensate. It is possible that a population of glycolytic and fermentation proteins stays more mobile in vir-condensates, whereas the remaining population of the co-opted cytosolic proteins is bound to the membrane-bound p33/p36, thus showing less mobility (Fig. 11). Indeed, BiFC experiments showed that a population of Pgk1 and Adh1 was bound to p33 within VROs (Fig S16) [38]. This model may explain the partial fluorescent recovery of the co-opted host proteins in VROs. Indeed, examples on the interactions between organellar membranes and membraneless condensates exist in the literature. Membrane surfaces and proteins in the membranes could organize condensates by restricting diffusion and promoting multivalent interactions between proteins. This lowers the critical concentration of proteins to undergo liquid-liquid phase separation [68].

The poor ATP production within VROs formed by the IDR mutants of TBSV p33, which are defective in vir-condensate formation, strongly indicates that the co-opted glycolytic and fermentation enzymes must be catalytically active in vir-condensates. This is further supported by additional data: (i) knocking down Adh1 or Pdc1 fermentation enzymes in plant leaves resulted in poor ATP level within VROs and significantly reduced TBSV replication [35]. These fermentation enzymes were poorly expressed in plant leaves, and highly expressed after TBSV infection. The fermentation enzymes are needed to regenerate NAD^+^ from pyruvate to allow continuous glycolysis within VROs. (ii) Knocking down several glycolytic enzymes (individually) in plant leaves also resulted in poor ATP level within VROs and significantly reduced TBSV replication [36, 37, 48]; (iii) knocking down the proteasomal Rpn11 key protein interaction ‘hub’, which is co-opted by TBSV p33 to facilitate the recruitment of glycolytic and fermentation enzymes into VROs, led to poor ATP level within VROs and significantly reduced TBSV replication (Fig S11) [38]; (iv) RavK protease-based destruction of the actin filaments, which are co-opted by TBSV to facilitate the recruitment of Rpn11 and glycolytic and fermentation enzymes into VROs, also resulted in poor ATP level within VROs and significantly reduced TBSV replication [38]. Chemical inhibition of mitochondrial TCA cycle in yeast, which mainly uses fermentation in high glucose media, did not inhibit TBSV replication [69], whereas inhibition of fermentation greatly inhibited TBSV replication in yeast [35]. All these data firmly confirmed that ATP must be produced within the TBSV VROs, therefore the co-opted glycolytic and fermentation enzymes must be catalytically active in vir-condensates of VROs in plants. The enrichment and high concentration of the glycolytic and fermentation enzymes in vir-condensates likely facilitates substrate channeling and efficient ATP production within VROs. Other types of biomolecular condensates are known to increase enzymatic activities. For example, enzymatic rates of protein SUMOylation can be increased up to 36-fold in a phase separated compartment [70], and glucose consumption and ATP production greatly increases in G (glycolytic) bodies [44].

We propose that the vir-condensate and the co-opted peroxisomes/mitochondria are held together within VROs by co-opted ER membranes [71], virus-induced membrane contact sites [27–30, 72] and the subverted actin filaments [62], forming entangled meshworks and resulting in rather stable and efficient VROs to produce vast numbers of progeny viral RNAs in a protected microenvironment (Fig 11). The presence of elaborate membranes and the actin meshwork likely causes the irregular shape of VROs and restricts the internal space within VROs. The gel-like behavior of vir-condensate might be facilitated by the complex meshworks of co-opted membranes and actin filaments, thus limiting the internal movement of larger proteins, but to a lesser extent of the movement of the smaller proteins within the VROs. The detailed molecular composition and the mechanism of vir-condensate formation will need further studies.

Although the vir-condensate and the VROs are unique structures within the virus-infected cells, it is known that glycolytic enzymes form condensates, called Glycolytic- or G-bodies, under various stress conditions [42, 44]. These G-bodies facilitate glucose utilization and rapid and increased ATP production via concentrating glycolytic enzymes to function as “metabolons” and to help cells survive the stress conditions. The basic theme of increased ATP production via sequestering glycolytic and fermentation enzymes in TBSV-induced vir-condensates might be analogous to that of G-bodies. It is important to note that vir-condensates associate with co-opted cellular membranes in VROs, whereas G-bodies are membraneless and much smaller in size than VROs. The overall molecular complexity of vir-condensates are likely higher than that of the G-bodies.

We also showed that the TBSV p33 or CIRV p36 replication proteins were capable of phase separation *in vitro*. FRAP assay showed no fluorescent recovery of p33C and p36C droplets, suggesting a gel-like condensate. In addition, partial resistance of droplets to 1,6-hexanediol suggest that electrostatic interactions contribute to p33 droplet formation. Interestingly, p33 and p36 replication proteins organized the partitioning of the co-opted cytosolic host proteins into p33/p36 droplets *in vitro*. This ability of the replication proteins might explain the key roles of p33/p36 in organization of vir-condensates *in planta*.

We showed that the extended IDR1 in TBSV p33 and CIRV p36 was critical for phase separation *in vitro*. Although the IDR1 alone supported droplet formation *in vitro*, those droplets were in liquid-like form. In contrast, the cytosol-exposed C-terminal portion of p33, which included both the IDR1 and the p33:p33/p92 interaction domain formed gel-like droplets *in vitro*, suggesting stronger interactions among the individual molecules within the droplets. Interestingly, the IDR1 was required for VRO biogenesis in *N. benthamiana* cells. Deletion of IDR1 in TBSV p33 and CIRV p36 inhibited the clustering of peroxisomes or mitochondria without interfering with the correct subcellular localization of these replication proteins. We defined a short, highly charged segment within IDR1, which was important for phase separation *in vitro*, for formation of functional vir-condensates, ATP production in VROs and it was essential for TBSV replication. These substitutions in IDR1 moderately affected the stability of p33 as shown in the replication assay. The possible reason is that p33-organized vir-condensates might protect the components of VROs from degradation through 26S proteasome or autophagy pathway. Interestingly, the IDR2 region was found to modulate droplet fluidity to some degree, allowing droplets to fuse a little longer forming larger droplets; or, the result may also indicate that the IDR2 region may contain ‘sticker’ regions [73], removal of which would result in larger droplets.

IDR sequences, which are frequently involved in multiple weak interactions, are known to facilitate phase separation in many different cellular proteins [74], and viral proteins [75–78]. The membraneless replication factories of several negative-strand RNA viruses consist of condensates formed by replication proteins and viral RNA in addition to co-opted host proteins [75, 77, 79]. Condensates formation is also required for various immune responses against infecting viruses [40, 79–81]. Condensate formation has been shown to affect plant +RNA virus movement [82]. Plant potyviruses induce RNA granules that are important for virus replication [83].

Overall, this paper demonstrates that tombusvirus VROs are complex structures containing distinct substructures with subverted membranes and vir-condensates. Because the vir-condensate is essential for VRO functions, interfering with vir-condensate formation might open up new antiviral approaches.

## Materials and methods

### Fluorescence recovery after photobleaching (FRAP)

VROs identified in agrobacterium-infiltrated *N. benthamiana* leaves were subjected to photobleaching. To avoid the movement of VROs along actin filaments, the leaves were treated with 10 μM Latrunculin B (Lat B) [Abcam] at least three hours before FRAP assay. Partial-, full- and half-bleaching were conducted on regions of interest (ROI) selected on VROs for 3 to 5 seconds with a 405 nm laser at 20-80% laser intensity, which was selected based on expression level of the given protein. Time-lapse images were taken every 20 sec within 2 min. More than 6 VROs enriched with co-opted different glycolytic and fermentation enzymes were subjected to FRAP assay and at least three independent plants were tested. As a control, cytosol-localized RFP-ADH1 was photobleached to compare the mobility of cytosolic proteins with proteins in gel-like vir-condensate. The images collected after FRAP assay were analyzed with the ROI manager tool of ImageJ software. The fluorescence intensities of ROIs were corrected using that of unbleached control regions and then normalized to the pre-bleached intensities of the ROIs for final quantification of fluorescence recovery efficiency. Fluorescence intensity dip for the unbleached half region in half-bleaching assay was detected and quantified following the method described by Muzzopappa et al [50]. The recovery curves and dip curves were plotted with Graphpad Prism9 software.

### *In vitro* droplet formation assay

For *in vitro* droplet formation, purified 5 μM TBSV eGFP-p33C and CIRV eGFP-p36C and their mutants were diluted in 40 μL reaction volume containing 30 mM Tris-HCl (pH 7.5), 150 mM NaCl and different concentrations of PEG8000 (10%, 5%, 2.5% or 0%) and then were incubated in 384 Well glass bottom plates (Cellvis) at room temperature and monitored for 10 to 40 min using an Olympus FV3000 confocal microscope. The co-droplet formation assay was performed as the droplet formation assay above, except additional 5 or 0.5 μM purified RFP-tagged host proteins were included. Purified His_6_-eGFP or His_6_-RFP were used as negative controls in the co-droplet formation assay. For FRAP assay, ROIs were selected on 8 individual droplets and photobleached for 3 sec with a 405 nm laser at 20% laser intensity using an Olympus FV3000 confocal microscope. Images were taken every 20 seconds within 2 min and image processing followed the same procedure as VRO-FRAP assay above. To monitor droplet fusion, time-lapse mode was used with an interval of 6.5 seconds for each scanning. To measure droplet size and quantify of droplet formed by different mutants, we followed the methods with some modification [80]. In brief, 8 randomly captured fields for each mutant with a field size of 42 μm×42 μm were analyzed with ImageJ software. The imported images were converted to greyscale (8-bit) at first and then the threshold for detection was set up as 18 to 255 for further analyzing particles (only the droplet size ≥0.2 μm^2^ were counted and analyzed).

### 1,6-Hexanediol treatment

To test whether vir-condensates exist in liquid-like or gel-like form, treatment with 1,6-Hexanediol, which can dissolve liquid-liquid phase separated assemblies through disrupting weak hydrophobic interactions between proteins, was used with 10 μg/ml Digitonin in *N. benthamiana* leaves agroinfiltrated to express the GFP, RFP or BFP-tagged viral and host proteins. Samples treated with only 10 μg/ml Digitonin were used as a negative control. To verify the function of 1,6-Hexanediol in plant leaves, the leaves of *N. benthamiana* transiently expressing GFP-AtDCP1, which labels the P body in plant cells, were used as a positive control. All the samples were monitored within one hour after infiltration using an Olympus FV3000 confocal microscope.

*In vitro* droplets formed by eGFP-p33C at 30 min in the presence of either 2.5% or 10% PEG8000 were treated with 10 μg/ml 1,6-Hexanediol, followed by monitoring droplets for 10 and 30 min by confocal microscope, as above.

### FRET-based measurement of ATP production in VROs

To measure ATP accumulation within p33 or mutants-induced VROs, the ATeam-based biosensor module was fused to the C-terminus of p33 and mutants, which were then expressed in plant cells via agroinfiltration as described [37]. Samples were checked at 2.5 days after agroinfiltration under the confocal microscopy. FRET values (YFP/CFP ratio) were obtained based on the quantification of CFP and Venus images using ImageJ software and the statistical data were then plotted with Graphpad Prism9 software.

## Acknowledgements

We thank Dr. Judit Pogany for critical reading of the manuscript. We also thank previous lab member Dr. M. Molho for providing confocal images.

## Supplement

Fig S1. **The co-opted Adh1 fermentation enzyme is present in condensate substructure in tombusvirus VROs.** (A-B) FRAP analysis demonstrates the partial fluorescence recovery of Adh1-BFP signal after photobleaching within a single VRO induced by either TBSV p33-RFP or CIRV p36-RFP in *N. benthamiana* cells. Note that the membrane-bound p33-RFP or p36-RFP signals were not recovered. Scale bars represent 10 μm. The graphs on the right show the extent of fluorescence recovery at a given time point. 6-to-8 VROs from three independent plants were tested. (C). FRAP analysis demonstrates the complete fluorescence recovery of RFP-Adh1 signal after photobleaching within the cytosolic area in uninfected *N. benthamiana* cells. The area circled by red was photobleached, whereas the area circled by green was not photobleached and used as a control. See further details in panel A.

**Fig S2. FRAP analysis of the co-opted glycolytic and fermentation enzymes in tombusvirus VROs.** The graphs show the extent of fluorescence recovery after photobleaching of the shown host factors within either TBSV or CIRV VROs in *N. benthamiana*. See further details in Fig S1.

**Fig S3. FRAP analysis of the co-opted glycolytic enzymes in tombusvirus VROs formed during virus infections.** FRAP analysis demonstrates the partial fluorescence recovery of the shown glycolytic enzymes after photobleaching within single VROs induced by either (A, C) TBSV or (F) CIRV infections in *N. benthamiana*. The clustered peroxisomes were marked with GFP-SKL peroxisomal marker, whereas the clustered mitochondria were visualized with GFP-Tim21. Scale bars represent 10 μm. The graphs on the right show the extent of fluorescence recovery of the given fluorescent host protein at a given time point within VROs. See further details in Fig S1.

**Fig S4. Rapid recovery of fluorescence signal of SUMO1 in condensate substructure in tombusvirus VROs.** Time course analysis of fluorescence recovery of the RFP-SUMO1 signal after photobleaching is shown within a single VRO induced by TBSV p33-BFP in a *N. benthamiana* cell. Note that the membrane-bound p33-BFP signal was not recovered. Scale bars represent 10 μm. The merged images of p33-BFP and RFP-SUMO1 are presented to show the extent of fluorescence recovery. 6-to-8 VROs from two independent plants were tested. The graph shows the extent of fluorescence recovery at a given time point.

**Fig S5. The effect of 1.6-hexanediol treatment on the vir-condensate in tombusvirus VROs.** (A-B) Leaves of *N. benthamiana* transiently co-expressed BFP-SKL peroxisomal marker and either GFP-Pdc1 or GFP-Rpn11. The plants were infected with TBSV to induce VROs. Then, 2.5 days later the leaves were infiltrated with 10% 1,6-hexanediol and imaged at 3, 15 and 30 min time points. An arrow points at the VRO consisting of clustered peroxisomes. The percentages at the right panel indicate little fluorescence reduction of GFP-Pdc1 or GFP-Rpn11 within TBSV-induced VROs due to 1,6-hexanediol treatment. (C-D) Similar experiments using 1,6-hexanediol-treated leaves of *N. benthamiana* transiently co-expressing the shown fluorescent proteins. Confocal imaging was performed at the 60 min time point. Scale bars represent 10 μm. (E) Control experiments to demonstrate the effect of 10% 1,6-hexanediol treatment on P-bodies formed in *N. benthamiana* transiently expressing GFP-Dcp1 marker. Note the almost complete disapperance of punctate P-bodies in the 1,6-hexanediol-treated cells. Scale bars represent 20 μm. Each experiment was repeated three times.

**Fig S6. The effect of 1,6-hexanediol treatment on droplets formed by the eGFP-p33C protein *in vitro*.** Confocal images of droplets formed by the eGFP-p33C in the presence of 2.5% or 10% PEG8000. 1,6-hexanediol treatment was performed after droplet formation, which lasted for 30 min. Scale bars represent 10 μm.

**Fig S7. Droplet formation by the tombusvirus replication protein mutants *in vitro*.** (A) Top: Scheme of the CIRV p36 replication protein with the known domains. The two deletion mutants are also shown. Bottom images: confocal images of droplets formed by CIRV eGFP-p36C, eGFP-IDR1 and the absence of droplets with eGFP-p36C-ΔIDR1. (B) The predicted conserved IDR1 and IDR2 regions in TBSV p33 and CIRV p36 are shown based on PONDR score. (C) Top images: confocal images of droplets formed by TBSV eGFP-p33C, eGFP-p33C-ΔIDR2 or eGFP-IDR1 in the order of A, B, C. Bottom left shows the droplet number, whereas bottom central panel shows the distribution of doplet size supported by the above proteins. Red box indicates droplet size of <1.0 μm^2^, green box represents 1.0-4.0 μm^2^, blue box represents the % of droplets of 4.0-8.0 μm^2^, whereas purple box indicates droplet size of >8.0 μm^2^. Bottom right panel shows a SDS-PAGE of the purified recombinant proteins. (D) Confocal imaging shows that the purified recombinant eGFP-p33C-ΔsIDR1 (schematically shown) produces mostly irregular aggregate forms in the presence of 10% PEG8000, whereas no droplets are observed in the presence of 2.5% PEG8000. Each experiment was repeated three times.

**Fig S8. The tombusvirus replication proteins organize partitioning of glycolytic and fermentation enzymes to droplets *in vitro*.** (A-B) Confocal images show partitioning of the given RFP-tagged plant or yeast proteins to droplets formed by purified recombinant TBSV eGFP-p33C or CIRV eGFP-p36C in the presence of 2.5% PEG8000. Control experiments (in a boxed area) show that eGFP did not form co-droplet with the given RFP-tagged plant or yeast proteins in the presence of 2.5% PEG8000 under the same conditions. Scale bars represent 10 μm. Each experiment was repeated three times.

**Fig S9. The tombusvirus replication proteins induce partitioning of the non-glycolytic Rpn11 to droplets *in vitro*.** (A-B) Confocal images show the partitioning of purified RFP-tagged plant or yeast Rpn11 proteins to droplets formed by purified recombinant TBSV eGFP-p33C or CIRV eGFP-p36C in the presence of 2.5% PEG8000. Control experiments show that RFP-Rpn11 and eGFP did not form co-droplet in the presence of 2.5% PEG8000 under the same conditions. Scale bars represent 10 μm. Each experiment was repeated three times.

**Fig S10. Droplet formation by the glycolytic and fermentation proteins *in vitro*.** Confocal images of droplets formed by the shown purified proteins in the presence of 10% PEG8000. Scale bars represent 10 μm.

**Fig S11. Reduced ATP production within the VROs in Rpn11 knockdown *N. benthamiana* cells.** (A) A scheme of FRET-based detection of ATP within the TBSV VRO. ATP biosensor (ATeam^YEMK^) was fused to p33 replication protein for targeting to VROs. See main text for additional details. (B) The proteasomal Rpn11 protein interaction ‘hub’ was knocked-down by VIGS in *N. benthamiana*. The more intense FRET signals are white and red (between 0.5 to 1.0 ratio), whereas the low FRET signals (0.1 and below) are light blue and dark blue. The VROs are marked with arrows. Three independent cells are shown.

**Fig S12. The tombusvirus VRO size does not significantly affect ATP production within the VROs.** The WT p33 induces various sized VROs consisting of different number of clustered peroxisomes in *N. benthamiana* cells. We measured ATP production in selected “small” (<10 clustered peroxisomes) or “large” (>10 clustered peroxisomes) VROs formed by p33 fused with an ATP biosensor (ATeam^YEMK^). The more intense FRET signals are white and red (between 0.5 to 1.0 ratio), whereas the low FRET signals (0.1 and below) are light blue and dark blue. We show the quantitative FRET values (obtained with ImageJ) for numbers of samples in the graph.

**Fig S13. Control experiments showing TBSV eGFP-p33-driven ER membrane enrichment within and around the VRO.** (A) Transient expression of GFP-HDEL or BFP-CTT was used to visualize the ER network in *N. benthamiana* cells in the absence (A-B) or present of eGFP-p33 (C). Arrows point at an individual VRO. See Fig 10 for further details. Scale bars represent 20 μm.

**Fig S14. Additional confocal images of the ER membrane meshwork formed around VROs** The VROs were marked by TBSV p33 or CIRV p36 and the vir-condensates with the shown RFP-tagged glycolytic proteins or Rpn11 in *N. benthamiana* cells infected with either TBSV or CIRV. See Fig 10 for further details.

**Fig S15. The co-opted actin filaments and the ER membranes form meshworks around the VROs.** Additional confocal images of the co-opted actin filaments and ER membrane meshwork formed around VROs marked by TBSV p33 or CIRV p36 in *N. benthamiana* cells infected with either TBSV or CIRV. See Fig 10 for further details.

**Fig S16. BiFC studies of the interaction between TBSV p33 and the host PGK1 or ADH1 within VROs in *N. benthamiana*.** Interaction between TBSV p33-cYFP replication protein and the nYFP-Pgk1 or Adh1 proteins was detected by BiFC. The merged images show the co-localization of RFP-SKL with the BiFC signal, indicating that the interaction between p33 replication protein and host proteins occurs in VROs in clustered peroxisomal membranes. See additional details in references [38, 48].

## References

1. Wang A. Dissecting the molecular network of virus-plant interactions: the complex roles of host factors. Annu Rev Phytopathol. 2015;53:45–66. Epub 2015/05/06. doi: 10.1146/annurev-phyto-080614-120001. PubMed PMID: 25938276.

2. Romero-Brey I, Bartenschlager R. Membranous replication factories induced by plus-strand RNA viruses. Viruses. 2014;6(7):2826–57. Epub 2014/07/24. doi: v6072826 [pii] 10.3390/v6072826. PubMed PMID: 25054883; PubMed Central PMCID: PMC4113795.

3. Nagy PD. Tombusvirus-Host Interactions: Co-Opted Evolutionarily Conserved Host Factors Take Center Court. Annu Rev Virol. 2016;3(1):491–515. Epub 2016/09/01. doi: 10.1146/annurev-virology-110615-042312. PubMed PMID: 27578441.

4. Jordan TX, Randall G. Flavivirus modulation of cellular metabolism. Curr Opin Virol. 2016;19:7–10. Epub 2016/06/10. doi: S1879-6257(16)30058-X [pii] 10.1016/j.coviro.2016.05.007. PubMed PMID: 27280383; PubMed Central PMCID: PMC5021554.

5. Shulla A, Randall G. (+) RNA virus replication compartments: a safe home for (most) viral replication. Curr Opin Microbiol. 2016;32:82–8. Epub 2016/06/03. doi: S1369-5274(16)30058-3 [pii] 10.1016/j.mib.2016.05.003. PubMed PMID: 27253151; PubMed Central PMCID: PMC4983521.

6. Syed GH, Amako Y, Siddiqui A. Hepatitis C virus hijacks host lipid metabolism. Trends Endocrinol Metab. 2010;21(1):33–40. Epub 2009/10/27. doi: S1043-2760(09)00147-7 [pii] 10.1016/j.tem.2009.07.005. PubMed PMID: 19854061; PubMed Central PMCID: PMC2818172.

7. den Boon JA, Diaz A, Ahlquist P. Cytoplasmic viral replication complexes. Cell Host Microbe. 2010;8(1):77–85. Epub 2010/07/20. doi: S1931-3128(10)00213-1 [pii] 10.1016/j.chom.2010.06.010. PubMed PMID: 20638644; PubMed Central PMCID: PMC2921950.

8. Altan-Bonnet N. Lipid Tales of Viral Replication and Transmission. Trends Cell Biol. 2017;27(3):201–13. Epub 2016/11/14. doi: S0962-8924(16)30153-2 [pii] 10.1016/j.tcb.2016.09.011. PubMed PMID: 27838086; PubMed Central PMCID: PMC5318230.

9. van der Schaar HM, Dorobantu CM, Albulescu L, Strating JR, van Kuppeveld FJ. Fat(al) attraction: Picornaviruses Usurp Lipid Transfer at Membrane Contact Sites to Create Replication Organelles. Trends Microbiol. 2016;24(7):535–46. Epub 2016/03/30. doi: S0966-842X(16)00054-8 [pii] 10.1016/j.tim.2016.02.017. PubMed PMID: 27020598.

10. Fernandez de Castro I, Tenorio R, Risco C. Virus assembly factories in a lipid world. Curr Opin Virol. 2016;18:20–6. Epub 2016/03/18. doi: S1879-6257(16)30013-X [pii] 10.1016/j.coviro.2016.02.009. PubMed PMID: 26985879.

11. Hyodo K, Okuno T. Hijacking of host cellular components as proviral factors by plant-infecting viruses. Adv Virus Res. 2020;107:37–86. Epub 2020/07/28. doi: 10.1016/bs.aivir.2020.04.002. PubMed PMID: 32711734.

12. Lazear HM, Diamond MS. New insights into innate immune restriction of West Nile virus infection. Curr Opin Virol. 2015;11:1–6. Epub 2015/01/03. doi: S1879-6257(14)00233-8 [pii] 10.1016/j.coviro.2014.12.001. PubMed PMID: 25554924; PubMed Central PMCID: PMC4456296.

13. Horner SM, Gale M, Jr. Regulation of hepatic innate immunity by hepatitis C virus. Nat Med. 2013;19(7):879–88. Epub 2013/07/10. doi: nm.3253 [pii] 10.1038/nm.3253. PubMed PMID: 23836238; PubMed Central PMCID: PMC4251871.

14. Schoggins JW, Rice CM. Innate immune responses to hepatitis C virus. Curr Top Microbiol Immunol. 2013;369:219–42. Epub 2013/03/07. doi: 10.1007/978-3-642-27340-7_9. PubMed PMID: 23463203.

15. Suthar MS, Diamond MS, Gale M, Jr. West Nile virus infection and immunity. Nat Rev Microbiol. 2013;11(2):115–28. Epub 2013/01/17. doi: nrmicro2950 [pii] 10.1038/nrmicro2950. PubMed PMID: 23321534.

16. Kovalev N, Inaba JI, Li Z, Nagy PD. The role of co-opted ESCRT proteins and lipid factors in protection of tombusviral double-stranded RNA replication intermediate against reconstituted RNAi in yeast. PLoS Pathog. 2017;13(7):e1006520. Epub 2017/08/02. doi: 10.1371/journal.ppat.1006520. PubMed PMID: 28759634; PubMed Central PMCID: PMCPMC5552349.

17. Garcia-Ruiz H. Susceptibility Genes to Plant Viruses. Viruses. 2018;10(9). Epub 2018/09/12. doi: 10.3390/v10090484. PubMed PMID: 30201857; PubMed Central PMCID: PMCPMC6164914.

18. Fernandez de Castro I, Fernandez JJ, Barajas D, Nagy PD, Risco C. Three-dimensional imaging of the intracellular assembly of a functional viral RNA replicase complex. J Cell Sci. 2017;130(1):260–8. Epub 2016/03/31. doi: 10.1242/jcs.181586. PubMed PMID: 27026525.

19. Barajas D, Martin IF, Pogany J, Risco C, Nagy PD. Noncanonical role for the host Vps4 AAA+ ATPase ESCRT protein in the formation of Tomato bushy stunt virus replicase. PLoS Pathog. 2014;10(4):e1004087. Epub 2014/04/26. doi: 10.1371/journal.ppat.1004087. PubMed PMID: 24763736; PubMed Central PMCID: PMCPMC3999190.

20. Nagy PD, Pogany J. The dependence of viral RNA replication on co-opted host factors. Nature Reviews Microbiology. 2012;10(2):137–49. doi: Doi 10.1038/Nrmicro2692. PubMed PMID: ISI:000299115000013.

21. White KA, Nagy PD. Advances in the molecular biology of tombusviruses: gene expression, genome replication, and recombination. Prog Nucleic Acid Res Mol Biol. 2004;78:187–226. Epub 2004/06/24. doi: 10.1016/S0079-6603(04)78005-8. PubMed PMID: 15210331.

22. Stork J, Kovalev N, Sasvari Z, Nagy PD. RNA chaperone activity of the tombusviral p33 replication protein facilitates initiation of RNA synthesis by the viral RdRp in vitro. Virology. 2011;409(2):338–47. Epub 2010/11/13. doi: 10.1016/j.virol.2010.10.015. PubMed PMID: 21071052; PubMed Central PMCID: PMCPMC7173327.

23. Pogany J, White KA, Nagy PD. Specific binding of tombusvirus replication protein p33 to an internal replication element in the viral RNA is essential for replication. J Virol. 2005;79(8):4859–69. Epub 2005/03/30. doi: 10.1128/JVI.79.8.4859-4869.2005. PubMed PMID: 15795271; PubMed Central PMCID: PMCPMC1069559.

24. Nagy PD, Feng Z. Tombusviruses orchestrate the host endomembrane system to create elaborate membranous replication organelles. Curr Opin Virol. 2021;48:30–41. Epub 20210410. doi: 10.1016/j.coviro.2021.03.007. PubMed PMID: 33845410.

25. Nagy PD. Co-opted membranes, lipids, and host proteins: what have we learned from tombusviruses? Curr Opin Virol. 2022;56:101258. Epub 20220923. doi: 10.1016/j.coviro.2022.101258. PubMed PMID: 36166851.

26. Nagy PD, Pogany J, Xu K. Cell-Free and Cell-Based Approaches to Explore the Roles of Host Membranes and Lipids in the Formation of Viral Replication Compartment Induced by Tombusviruses. Viruses. 2016;8(3). Epub 2016/03/08. doi: v8030068 [pii] 10.3390/v8030068. PubMed PMID: 26950140.

27. Barajas D, Xu K, de Castro Martin IF, Sasvari Z, Brandizzi F, Risco C, et al. Co-opted oxysterol-binding ORP and VAP proteins channel sterols to RNA virus replication sites via membrane contact sites. PLoS Pathog. 2014;10(10):e1004388. Epub 2014/10/21. doi: 10.1371/journal.ppat.1004388. PubMed PMID: 25329172; PubMed Central PMCID: PMCPMC4199759.

28. Nagy PD, Strating JR, van Kuppeveld FJ. Building Viral Replication Organelles: Close Encounters of the Membrane Types. PLoS Pathog. 2016;12(10):e1005912. Epub 2016/10/28. doi: 10.1371/journal.ppat.1005912. PubMed PMID: 27788266; PubMed Central PMCID: PMCPMC5082816.

29. Sasvari Z, Lin W, Inaba JI, Xu K, Kovalev N, Nagy PD. Co-opted Cellular Sac1 Lipid Phosphatase and PI(4)P Phosphoinositide Are Key Host Factors during the Biogenesis of the Tombusvirus Replication Compartment. J Virol. 2020;94(12). Epub 2020/04/10. doi: 10.1128/JVI.01979-19. PubMed PMID: 32269127; PubMed Central PMCID: PMCPMC7307105.

30. Lin W, Feng Z, Prasanth KR, Liu Y, Nagy PD. Dynamic interplay between the co-opted Fis1 mitochondrial fission protein and membrane contact site proteins in supporting tombusvirus replication. PLoS Pathog. 2021;17(3):e1009423. Epub 20210316. doi: 10.1371/journal.ppat.1009423. PubMed PMID: 33725015; PubMed Central PMCID: PMCPMC7997005.

31. Feng Z, Inaba JI, Nagy PD. The retromer is co-opted to deliver lipid enzymes for the biogenesis of lipid-enriched tombusviral replication organelles. Proc Natl Acad Sci U S A. 2021;118(1). Epub 2020/12/31. doi: 10.1073/pnas.2016066118. PubMed PMID: 33376201; PubMed Central PMCID: PMCPMC7817191.

32. Inaba JI, Xu K, Kovalev N, Ramanathan H, Roy CR, Lindenbach BD, et al. Screening Legionella effectors for antiviral effects reveals Rab1 GTPase as a proviral factor coopted for tombusvirus replication. Proc Natl Acad Sci U S A. 2019;116(43):21739–47. Epub 2019/10/09. doi: 10.1073/pnas.1911108116. PubMed PMID: 31591191; PubMed Central PMCID: PMCPMC6815150.

33. Xu K, Nagy PD. Enrichment of Phosphatidylethanolamine in Viral Replication Compartments via Co-opting the Endosomal Rab5 Small GTPase by a Positive-Strand RNA Virus. PLoS Biol. 2016;14(10):e2000128. Epub 2016/10/21. doi: 10.1371/journal.pbio.2000128. PubMed PMID: 27760128; PubMed Central PMCID: PMCPMC5070881.

34. Nagy PD, Lin W. Taking over Cellular Energy-Metabolism for TBSV Replication: The High ATP Requirement of an RNA Virus within the Viral Replication Organelle. Viruses. 2020;12(1). Epub 2020/01/18. doi: 10.3390/v12010056. PubMed PMID: 31947719; PubMed Central PMCID: PMCPMC7019945.

35. Lin W, Liu Y, Molho M, Zhang S, Wang L, Xie L, et al. Co-opting the fermentation pathway for tombusvirus replication: Compartmentalization of cellular metabolic pathways for rapid ATP generation. PLoS Pathog. 2019;15(10):e1008092. Epub 2019/10/28. doi: 10.1371/journal.ppat.1008092. PubMed PMID: 31648290; PubMed Central PMCID: PMCPMC6830812.

36. Prasanth KR, Chuang C, Nagy PD. Co-opting ATP-generating glycolytic enzyme PGK1 phosphoglycerate kinase facilitates the assembly of viral replicase complexes. PLoS Pathog. 2017;13(10):e1006689. Epub 2017/10/24. doi: 10.1371/journal.ppat.1006689. PubMed PMID: 29059239; PubMed Central PMCID: PMCPMC5695612.

37. Chuang C, Prasanth KR, Nagy PD. The Glycolytic Pyruvate Kinase Is Recruited Directly into the Viral Replicase Complex to Generate ATP for RNA Synthesis. Cell Host Microbe. 2017;22(5):639–52 e7. Epub 2017/11/07. doi: 10.1016/j.chom.2017.10.004. PubMed PMID: 29107644.

38. Molho M, Lin W, Nagy PD. A novel viral strategy for host factor recruitment: The co-opted proteasomal Rpn11 protein interaction hub in cooperation with subverted actin filaments are targeted to deliver cytosolic host factors for viral replication. PLoS Pathog. 2021;17(6):e1009680. Epub 20210623. doi: 10.1371/journal.ppat.1009680. PubMed PMID: 34161398; PubMed Central PMCID: PMCPMC8260003.

39. Alberti S, Hyman AA. Biomolecular condensates at the nexus of cellular stress, protein aggregation disease and ageing. Nat Rev Mol Cell Biol. 2021;22(3):196–213. Epub 2021/01/30. doi: 10.1038/s41580-020-00326-6. PubMed PMID: 33510441.

40. Zavaliev R, Mohan R, Chen T, Dong X. Formation of NPR1 Condensates Promotes Cell Survival during the Plant Immune Response. Cell. 2020;182(5):1093–108 e18. Epub 2020/08/19. doi: 10.1016/j.cell.2020.07.016. PubMed PMID: 32810437; PubMed Central PMCID: PMCPMC7484032.

41. Banani SF, Lee HO, Hyman AA, Rosen MK. Biomolecular condensates: organizers of cellular biochemistry. Nat Rev Mol Cell Biol. 2017;18(5):285–98. Epub 2017/02/23. doi: 10.1038/nrm.2017.7. PubMed PMID: 28225081; PubMed Central PMCID: PMCPMC7434221.

42. Fuller GG, Kim JK. Compartmentalization and metabolic regulation of glycolysis. J Cell Sci. 2021;134(20). Epub 2021/10/21. doi: 10.1242/jcs.258469. PubMed PMID: 34668544; PubMed Central PMCID: PMCPMC8572002.

43. Fuller GG, Han T, Freeberg MA, Moresco JJ, Ghanbari Niaki A, Roach NP, et al. RNA promotes phase separation of glycolysis enzymes into yeast G bodies in hypoxia. Elife. 2020;9. Epub 2020/04/17. doi: 10.7554/eLife.48480. PubMed PMID: 32298230; PubMed Central PMCID: PMCPMC7162659.

44. Jin M, Fuller GG, Han T, Yao Y, Alessi AF, Freeberg MA, et al. Glycolytic Enzymes Coalesce in G Bodies under Hypoxic Stress. Cell Rep. 2017;20(4):895–908. Epub 2017/07/27. doi: 10.1016/j.celrep.2017.06.082. PubMed PMID: 28746874; PubMed Central PMCID: PMCPMC5586494.

45. Milovanovic D, Wu Y, Bian X, De Camilli P. A liquid phase of synapsin and lipid vesicles. Science. 2018;361(6402):604–7. Epub 2018/07/07. doi: 10.1126/science.aat5671. PubMed PMID: 29976799; PubMed Central PMCID: PMCPMC6191856.

46. Gao Y, Zhu Y, Wang H, Cheng Y, Zhao D, Sun Q, et al. Lipid-mediated phase separation of AGO proteins on the ER controls nascent-peptide ubiquitination. Mol Cell. 2022;82(7):1313–28 e8. Epub 20220323. doi: 10.1016/j.molcel.2022.02.035. PubMed PMID: 35325613.

47. Nagy PD. Host protein chaperones, RNA helicases and the ubiquitin network highlight the arms race for resources between tombusviruses and their hosts. Adv Virus Res. 2020;107:133–58. Epub 2020/07/28. doi: 10.1016/bs.aivir.2020.06.006. PubMed PMID: 32711728; PubMed Central PMCID: PMCPMC7342006.

48. Molho M, Chuang C, Nagy PD. Co-opting of nonATP-generating glycolytic enzymes for TBSV replication. Virology. 2021;559:15–29. Epub 20210322. doi: 10.1016/j.virol.2021.03.011. PubMed PMID: 33799077.

49. McSwiggen DT, Mir M, Darzacq X, Tjian R. Evaluating phase separation in live cells: diagnosis, caveats, and functional consequences. Genes Dev. 2019;33(23-24):1619–34. Epub 20191008. doi: 10.1101/gad.331520.119. PubMed PMID: 31594803; PubMed Central PMCID: PMCPMC6942051.

50. Muzzopappa F, Hummert J, Anfossi M, Tashev SA, Herten DP, Erdel F. Detecting and quantifying liquid-liquid phase separation in living cells by model-free calibrated half-bleaching. Nat Commun. 2022;13(1):7787. Epub 20221216. doi: 10.1038/s41467-022-35430-y. PubMed PMID: 36526633; PubMed Central PMCID: PMCPMC9758202.

51. Prasanth KR, Barajas D, Nagy PD. The proteasomal Rpn11 metalloprotease suppresses tombusvirus RNA recombination and promotes viral replication via facilitating assembly of the viral replicase complex. J Virol. 2015;89(5):2750–63. Epub 2014/12/30. doi: 10.1128/JVI.02620-14. PubMed PMID: 25540361; PubMed Central PMCID: PMCPMC4325744.

52. Feng Z, Kovalev N, Nagy PD. Multifunctional role of the co-opted Cdc48 AAA+ ATPase in tombusvirus replication. Virology. 2022;576:1–17. Epub 20220913. doi: 10.1016/j.virol.2022.08.004. PubMed PMID: 36126429.

53. Yu X, Li B, Jang GJ, Jiang S, Jiang D, Jang JC, et al. Orchestration of Processing Body Dynamics and mRNA Decay in Arabidopsis Immunity. Cell Rep. 2019;28(8):2194–205 e6. doi: 10.1016/j.celrep.2019.07.054. PubMed PMID: 31433992; PubMed Central PMCID: PMCPMC6716526.

54. Pak CW, Kosno M, Holehouse AS, Padrick SB, Mittal A, Ali R, et al. Sequence Determinants of Intracellular Phase Separation by Complex Coacervation of a Disordered Protein. Mol Cell. 2016;63(1):72–85. doi: 10.1016/j.molcel.2016.05.042. PubMed PMID: 27392146; PubMed Central PMCID: PMCPMC4973464.

55. Panavas T, Hawkins CM, Panaviene Z, Nagy PD. The role of the p33:p33/p92 interaction domain in RNA replication and intracellular localization of p33 and p92 proteins of Cucumber necrosis tombusvirus. Virology. 2005;338(1):81–95. Epub 2005/06/07. doi: 10.1016/j.virol.2005.04.025. PubMed PMID: 15936051.

56. Gunawardene CD, Donaldson LW, White KA. Tombusvirus polymerase: Structure and function. Virus Res. 2017;234:74–86. Epub 2017/01/24. doi: S0168-1702(16)30702-X [pii] 10.1016/j.virusres.2017.01.012. PubMed PMID: 28111194.

57. Rajendran KS, Nagy PD. Characterization of the RNA-binding domains in the replicase proteins of tomato bushy stunt virus. J Virol. 2003;77(17):9244–58. Epub 2003/08/14. doi: 10.1128/jvi.77.17.9244-9258.2003. PubMed PMID: 12915540; PubMed Central PMCID: PMCPMC187376.

58. McCartney AW, Greenwood JS, Fabian MR, White KA, Mullen RT. Localization of the tomato bushy stunt virus replication protein p33 reveals a peroxisome-to-endoplasmic reticulum sorting pathway. Plant Cell. 2005;17(12):3513–31. Epub 2005/11/15. doi: tpc.105.036350 [pii] 10.1105/tpc.105.036350. PubMed PMID: 16284309; PubMed Central PMCID: PMC1315385.

59. Schuster BS, Dignon GL, Tang WS, Kelley FM, Ranganath AK, Jahnke CN, et al. Identifying sequence perturbations to an intrinsically disordered protein that determine its phase-separation behavior. Proc Natl Acad Sci U S A. 2020;117(21):11421–31. Epub 20200511. doi: 10.1073/pnas.2000223117. PubMed PMID: 32393642; PubMed Central PMCID: PMCPMC7261017.

60. Imamura H, Nhat KP, Togawa H, Saito K, Iino R, Kato-Yamada Y, et al. Visualization of ATP levels inside single living cells with fluorescence resonance energy transfer-based genetically encoded indicators. Proc Natl Acad Sci U S A. 2009;106(37):15651–6. Epub 20090831. doi: 10.1073/pnas.0904764106. PubMed PMID: 19720993; PubMed Central PMCID: PMCPMC2735558.

61. Nagano M, Ueda H, Fukao Y, Kawai-Yamada M, Hara-Nishimura I. Generation of Arabidopsis lines with a red fluorescent marker for endoplasmic reticulum using a tail-anchored protein cytochrome b(5) -B. Plant Signal Behav. 2020;15(9):1790196. Epub 20200707. doi: 10.1080/15592324.2020.1790196. PubMed PMID: 32633191; PubMed Central PMCID: PMCPMC8550181.

62. Nawaz-ul-Rehman MS, Prasanth KR, Xu K, Sasvari Z, Kovalev N, de Castro Martin IF, et al. Viral Replication Protein Inhibits Cellular Cofilin Actin Depolymerization Factor to Regulate the Actin Network and Promote Viral Replicase Assembly. PLoS Pathog. 2016;12(2):e1005440. Epub 2016/02/11. doi: 10.1371/journal.ppat.1005440. PubMed PMID: 26863541; PubMed Central PMCID: PMCPMC4749184.

63. Molho M, Zhu S, Nagy PD. Race against Time between the Virus and Host: Actin-Assisted Rapid Biogenesis of Replication Organelles is Used by TBSV to Limit the Recruitment of Cellular Restriction Factors. J Virol. 2022:e0016821. Epub 20220531. doi: 10.1128/jvi.00168-21. PubMed PMID: 35638821.

64. Neufeldt CJ, Cortese M, Acosta EG, Bartenschlager R. Rewiring cellular networks by members of the Flaviviridae family. Nat Rev Microbiol. 2018;16(3):125–42. Epub 2018/02/13. doi: nrmicro.2017.170 [pii] 10.1038/nrmicro.2017.170. PubMed PMID: 29430005; PubMed Central PMCID: PMC7097628.

65. de Wilde AH, Snijder EJ, Kikkert M, van Hemert MJ. Host Factors in Coronavirus Replication. Curr Top Microbiol Immunol. 2018;419:1–42. Epub 2017/06/24. doi: 10.1007/82_2017_25. PubMed PMID: 28643204; PubMed Central PMCID: PMC7119980.

66. Paul D, Bartenschlager R. Flaviviridae Replication Organelles: Oh, What a Tangled Web We Weave. Annu Rev Virol. 2015;2(1):289–310. Epub 2016/03/10. doi: 10.1146/annurev-virology-100114-055007. PubMed PMID: 26958917.

67. Rademacher DJ, Bello AI, May JP. CASC3 Biomolecular Condensates Restrict Turnip Crinkle Virus by Limiting Host Factor Availability. J Mol Biol. 2023:167956. Epub 20230113. doi: 10.1016/j.jmb.2023.167956. PubMed PMID: 36642157; PubMed Central PMCID: PMCPMC10338645.

68. Zhao YG, Zhang H. Phase Separation in Membrane Biology: The Interplay between Membrane-Bound Organelles and Membraneless Condensates. Dev Cell. 2020;55(1):30–44. Epub 20200728. doi: 10.1016/j.devcel.2020.06.033. PubMed PMID: 32726575.

69. Inaba JI, Nagy PD. Tombusvirus RNA replication depends on the TOR pathway in yeast and plants. Virology. 2018;519:207–22. Epub 2018/05/08. doi: 10.1016/j.virol.2018.04.010. PubMed PMID: 29734044.

70. Peeples W, Rosen MK. Mechanistic dissection of increased enzymatic rate in a phase-separated compartment. Nat Chem Biol. 2021;17(6):693–702. Epub 20210525. doi: 10.1038/s41589-021-00801-x. PubMed PMID: 34035521; PubMed Central PMCID: PMCPMC8635274.

71. Sasvari Z, Kovalev N, Gonzalez PA, Xu K, Nagy PD. Assembly-hub function of ER-localized SNARE proteins in biogenesis of tombusvirus replication compartment. PLoS Pathog. 2018;14(5):e1007028. Epub 2018/05/11. doi: 10.1371/journal.ppat.1007028. PubMed PMID: 29746582; PubMed Central PMCID: PMCPMC5963807.

72. Kang Y, Lin W, Liu Y, Nagy PD. Key tethering function of Atg11 autophagy scaffold protein in formation of virus-induced membrane contact sites during tombusvirus replication. Virology. 2022;572:1–16. Epub 20220429. doi: 10.1016/j.virol.2022.04.007. PubMed PMID: 35533414.

73. Dar F, Pappu R. Restricting the sizes of condensates. Elife. 2020;9. Epub 20200714. doi: 10.7554/eLife.59663. PubMed PMID: 32662769; PubMed Central PMCID: PMCPMC7360362.

74. Borcherds W, Bremer A, Borgia MB, Mittag T. How do intrinsically disordered protein regions encode a driving force for liquid-liquid phase separation? Curr Opin Struct Biol. 2021;67:41–50. Epub 20201015. doi: 10.1016/j.sbi.2020.09.004. PubMed PMID: 33069007; PubMed Central PMCID: PMCPMC8044266.

75. Lopez N, Camporeale G, Salgueiro M, Borkosky SS, Visentin A, Peralta-Martinez R, et al. Deconstructing virus condensation. PLoS Pathog. 2021;17(10):e1009926. Epub 20211014. doi: 10.1371/journal.ppat.1009926. PubMed PMID: 34648608; PubMed Central PMCID: PMCPMC8516229.

76. Rahman SK, Ampah KK, Roy P. Role of NS2 specific RNA binding and phosphorylation in liquid-liquid phase separation and virus assembly. Nucleic Acids Res. 2022;50(19):11273–84. doi: 10.1093/nar/gkac904. PubMed PMID: 36259663; PubMed Central PMCID: PMCPMC9638936.

77. Wu C, Holehouse AS, Leung DW, Amarasinghe GK, Dutch RE. Liquid Phase Partitioning in Virus Replication: Observations and Opportunities. Annu Rev Virol. 2022;9(1):285–306. Epub 20220616. doi: 10.1146/annurev-virology-093020-013659. PubMed PMID: 35709511.

78. Ambroggio EE, Costa Navarro GS, Perez Socas LB, Bagatolli LA, Gamarnik AV. Dengue and Zika virus capsid proteins bind to membranes and self-assemble into liquid droplets with nucleic acids. J Biol Chem. 2021;297(3):101059. Epub 20210808. doi: 10.1016/j.jbc.2021.101059. PubMed PMID: 34375636; PubMed Central PMCID: PMCPMC8397897.

79. Brocca S, Grandori R, Longhi S, Uversky V. Liquid-Liquid Phase Separation by Intrinsically Disordered Protein Regions of Viruses: Roles in Viral Life Cycle and Control of Virus-Host Interactions. Int J Mol Sci. 2020;21(23). Epub 20201128. doi: 10.3390/ijms21239045. PubMed PMID: 33260713; PubMed Central PMCID: PMCPMC7730420.

80. Fang XD, Gao Q, Zang Y, Qiao JH, Gao DM, Xu WY, et al. Host casein kinase 1-mediated phosphorylation modulates phase separation of a rhabdovirus phosphoprotein and virus infection. Elife. 2022;11. Epub 20220222. doi: 10.7554/eLife.74884. PubMed PMID: 35191833; PubMed Central PMCID: PMCPMC8887900.

81. Shen C, Li R, Negro R, Cheng J, Vora SM, Fu TM, et al. Phase separation drives RNA virus-induced activation of the NLRP6 inflammasome. Cell. 2021;184(23):5759–74 e20. Epub 20211021. doi: 10.1016/j.cell.2021.09.032. PubMed PMID: 34678144; PubMed Central PMCID: PMCPMC8643277.

82. Brown SL, Garrison DJ, May JP. Phase separation of a plant virus movement protein and cellular factors support virus-host interactions. PLoS Pathog. 2021;17(9):e1009622. Epub 20210920. doi: 10.1371/journal.ppat.1009622. PubMed PMID: 34543360; PubMed Central PMCID: PMCPMC8483311.

83. Hafren A, Lohmus A, Makinen K. Formation of Potato Virus A-Induced RNA Granules and Viral Translation Are Interrelated Processes Required for Optimal Virus Accumulation. PLoS Pathog. 2015;11(12):e1005314. Epub 20151207. doi: 10.1371/journal.ppat.1005314. PubMed PMID: 26641460; PubMed Central PMCID: PMCPMC4671561.

