## supplemental figure for "Co-opted cytosolic host proteins form unique condensate substructures within the membranous tombusviral replication organelles"

**Supplement**

**Materials and methods**

**Plant materials, yeast strain and plasmids.** Wild type *Nicotiana benthamiana* plants were potted in soil and grown in greenhouse for three to four weeks. Then, similar sized plants were moved to the growth room and grown under a 16-h-light/8-h-dark cycle and a constant temperature at 25°C for one week before agroinfiltration. *S. cerevisiae* strain BY4741 (MATa his3Δ1 leu2Δ0 met15Δ0 ura3Δ0) was purchased from Open Biosystems and stored in -80 °C refrigerator. Plasmids used in this study and the primers used for plasmid construction are listed in Table S1. TBSV p33 and CIRV p36 mutants used in this study were sequenced by ACGT, Inc (USA).

**Agroinfiltration and confocal microscopy imaging.** To check virus replication organelles (VRO), TBSV p33 or CIRV p36 replication proteins and glycolytic and fermentation enzymes, plus Rpn11 proteasomal protein were tagged with fluorescent proteins. *N. benthamiana* leaves were agroinfiltrated in different combinations using agrobacteria ($OD_{600}$ 0.3 for each). The fluorescence signals were checked and captured with an Olympus FV3000 confocal microscope under a 60× silicon objective at 2.5 to 3 days post agroinfiltration (dpai). To check TBSV or CIRV infection-induced VROs, peroxisomal marker BFP-SKL or GFP-SKL or mitochondrial marker CoxIV-BFP or GFP-AtTim21 were co-expressed with fluorescent protein-tagged glycolytic and fermentation enzymes in *N. benthamiana* leaves via agrobacterium. Then, 16 h later, the agroinfiltrated leaves were inoculated with TBSV or CIRV sap. Endoplasmic reticulum was visualized either with a ER-targeting transmembrane domain of *Arabidopsis* cytochrome b5-B protein tagged with BFP or GFP or a ER-lumen-targeted fluorescent protein with both an N-terminal signal peptide (SP) and a C-terminal ER retention tetrapeptide (GFP-HDEL) (1). Actin filaments were visualized using LifeAct-RFP, which features a 17-amino-acid peptide specifically binding to filamentous actin (F-actin) structures in eukaryotic cells (2). The images were captured at 3 days post sap inoculation (dpi). BFP, GFP and RFP were excited at 405, 488 and 561 nm, respectively, and detected at 430-470, 495-535, and 560-600 nm using an Olympus FV3000 confocal microscope.

**Protein purification from *E. Coli* BL21**. All the recombinant proteins used in this study were purified from *E. Coli* BL21(DE3) (3). Briefly, all pET-30a based expression vectors were transformed into *E. Coli* BL21 competent cells. A single colony for each construct was picked up and grown in 3 mL MB broth containing 100 µg/mL kanamycin and 34 µg/mL chloramphenicol at 37°C overnight. Then the pre-growing culture was diluted 200-fold in 300 mL MB broth containing 100 µg/mL kanamycin and 34 µg/mL chloramphenicol and grown at 37°C for about 3.5 h until the $OD_{600}$ reached ~0.7. After 10 min of incubation on ice, 0.3 mM isopropyl β-D-1-thiogalactopyranoside (IPTG) was added into each culture to induce protein expression at 14°C for 18 h. The shaker speed for growing bacterium was always kept at 250 rpm. Bacterium cells were then harvested into three aliquots and each aliquot was suspended in 5 ml extraction buffer (30 mM Tris-HCl pH 7.5, 500 mM NaCl and 20 mM imidazole) for further sonication. The sonicated cells were centrifuged at 4°C for 20 min with max speed to obtain supernatant fraction, which was immediately incubated with 300 µL ProBond™ Nickel-Chelating Resin (Invitrogen) equilibrated with extraction buffer in a column at 4°C for 1 h with a rotator speed at 5 rpm. After draining by gravity and washing with buffer (30 mM Tris-HCl pH 7.5, 500 mM NaCl and 40 mM imidazole), the purified proteins were obtained with 500 µL elution buffer (30 mM Tris-HCl pH 7.5, 500 mM NaCl, and 250 mM imidazole) and 20 µL of each protein was aliquoted and stored at -80°C. Note that TBSV eGFP-p33C and CIRV eGFP-p36C proteins are prone to aggregation in high salt extraction buffer (500 mM NaCl). Therefore, physiological salt concentration (150 mM NaCl) was applied in all the purification process. The concentration of all the proteins purified in this study was measured using a standard curve. The purified $His_6$-tagged proteins were analyzed by SDS-PAGE electrophoresis followed by Coomassie staining to confirm their purity.

**Virus replication assay in yeast.** To test if the TBSV p33-IDR1m1 could support virus replication, the plasmid-based expression of p33, p92 and DI-72 replicon RNA was applied in yeast as described (4, 5). Briefly, plasmids LpGAD-Gal-Hisp92, UpYES-EV and HpESC-Gal-Hisp33 (or His33-IDR1m1)/Gal-DI72 were co-transformed into yeast strain BY4741. Transformed yeast cells were pre-grown in 2 ml SC-ULH$^{-}$ medium supplemented with 2% glucose for 16 h at 23°C. Then, yeast cultures were resuspended in SC-ULH$^{-}$ medium supplemented with 2% galactose and grown for 48 h at 23°C. We applied the standard RNA and protein analysis for yeast cells as previously described (5).

#### References

1. M. Nagano, H. Ueda, Y. Fukao, M. Kawai-Yamada, I. Hara-Nishimura, Generation of Arabidopsis lines with a red fluorescent marker for endoplasmic reticulum using a tail-anchored protein cytochrome b5 -B. *Plant Signal Behav* **15**, 1790196 (2020).
2. J. Riedl *et al.*, Lifeact: a versatile marker to visualize F-actin. *Nat Methods* **5**, 605-607 (2008).

3. K. S. Rajendran, P. D. Nagy, Characterization of the RNA-binding domains in the replicase proteins of tomato bushy stunt virus. *J Virol* **77**, 9244-9258 (2003).
4. T. Panavas, E. Serviene, J. Pogany, P. D. Nagy, Genome-wide screens for identification of host factors in viral replication. *Methods Mol Biol* **451**, 615-624 (2008).
5. T. Panavas, E. Serviene, J. Brasher, P. D. Nagy, Yeast genome-wide screen reveals dissimilar sets of host genes affecting replication of RNA viruses. *Proc Natl Acad Sci U S A* **102**, 7326-7331 (2005).

FIG S1

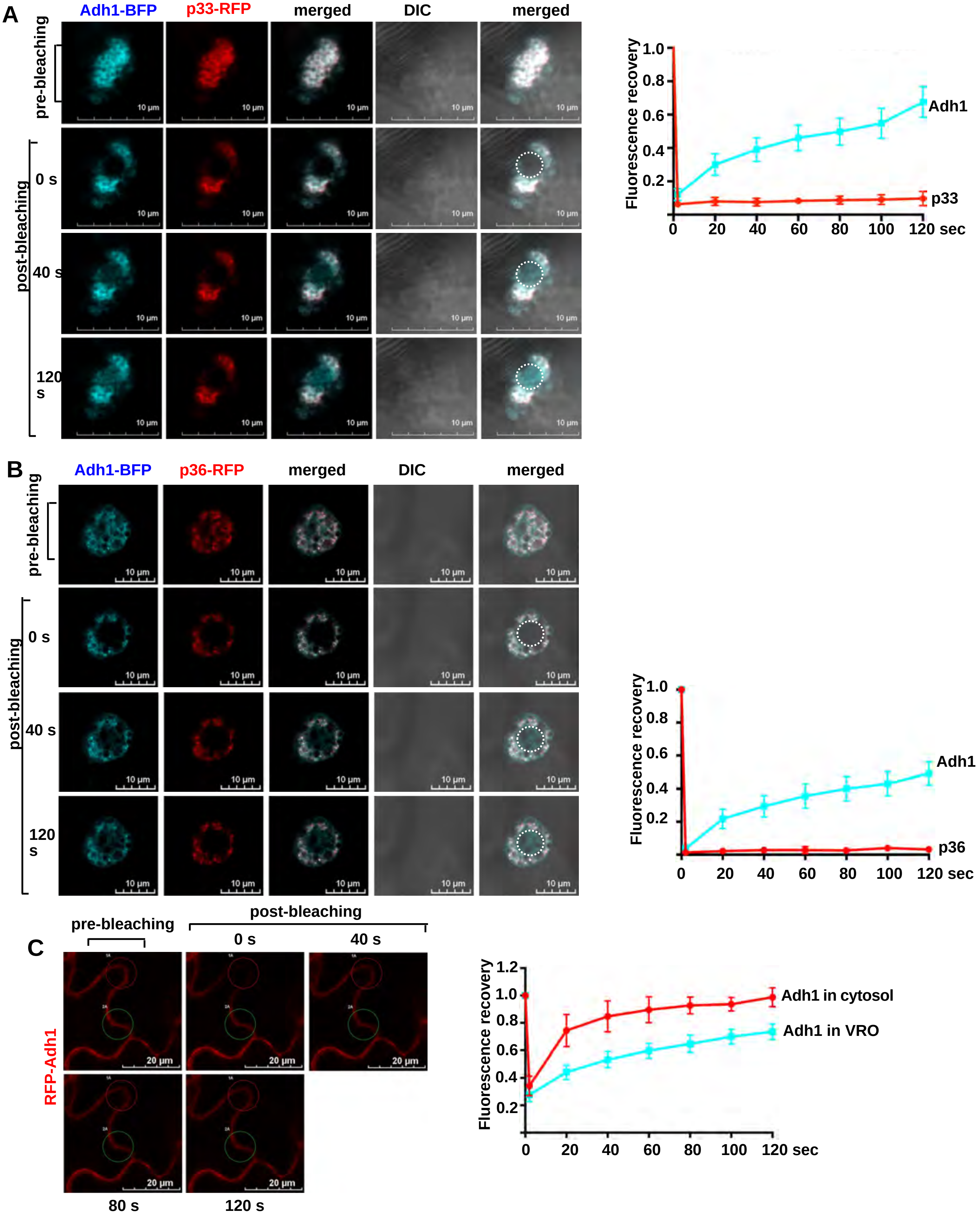
A
Adh1-BFP
p33-RFP
merged
DIC
merged
pre-bleaching
post-bleaching
0 s
40 s
120 s
10 µm
Fluorescence recovery
1.0
0.8
0.6
0.4
0.2
0
20
40
60
80
100
120 sec
Adh1
p33
B
Adh1-BFP
p36-RFP
merged
DIC
merged
pre-bleaching
post-bleaching
0 s
40 s
120 s
10 µm
Fluorescence recovery
Adh1
p36
C
pre-bleaching
post-bleaching
0 s
40 s
80 s
120 s
RFP-Adh1
20 µm
Fluorescence recovery
1.2
Adh1 in cytosol
Adh1 in VRO

#### FIG S2

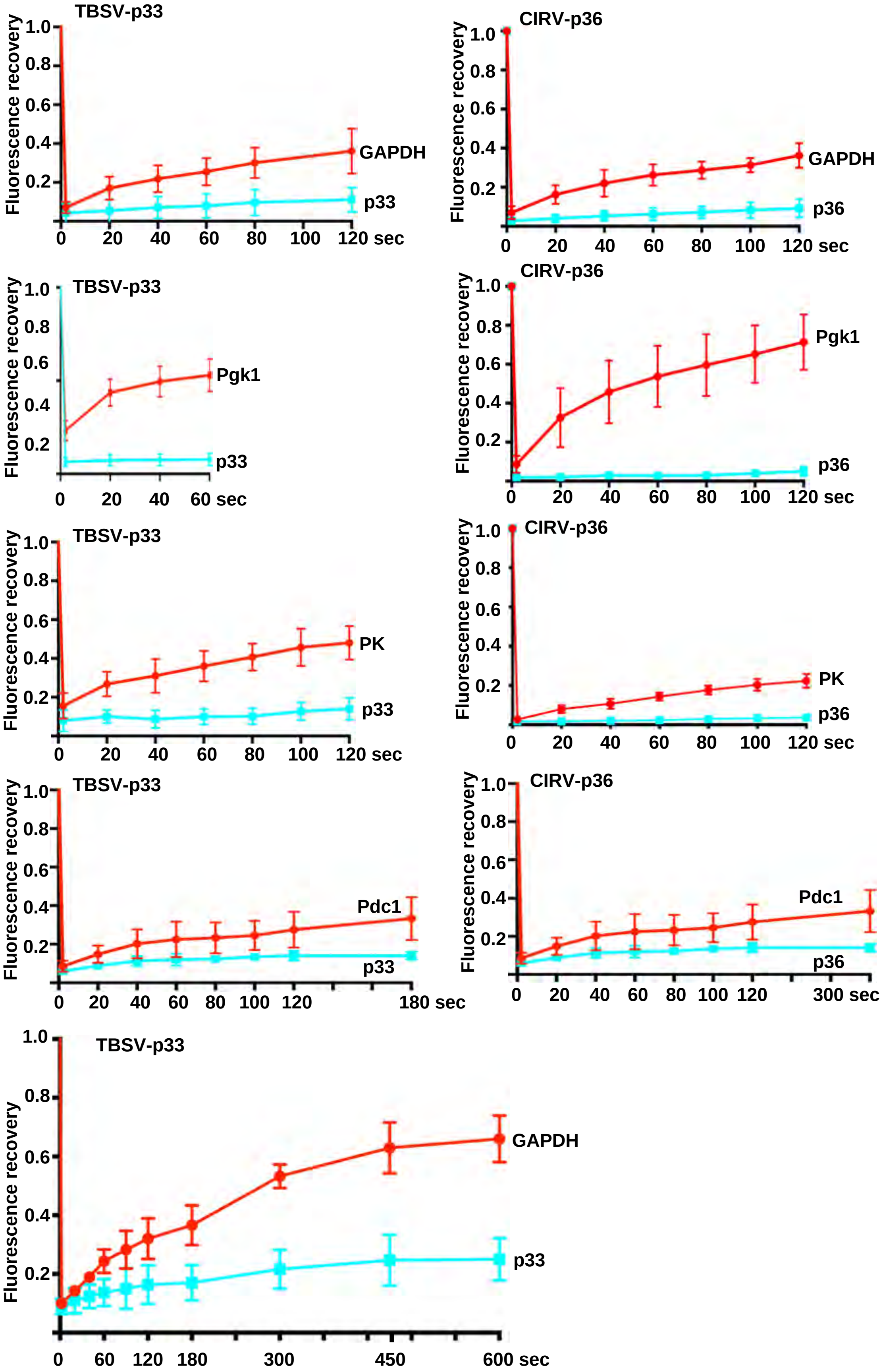

TBSV-p33
CIRV-p36
Fluorescence recovery
1.0
0.8
0.6
0.4
0.2
GAPDH
p33
p36
Pgk1
PK
Pdc1
0
20
40
60
80
100
120 sec
60 sec
180 sec
300 sec
120
180
300
450
600 sec

#### FIG S3

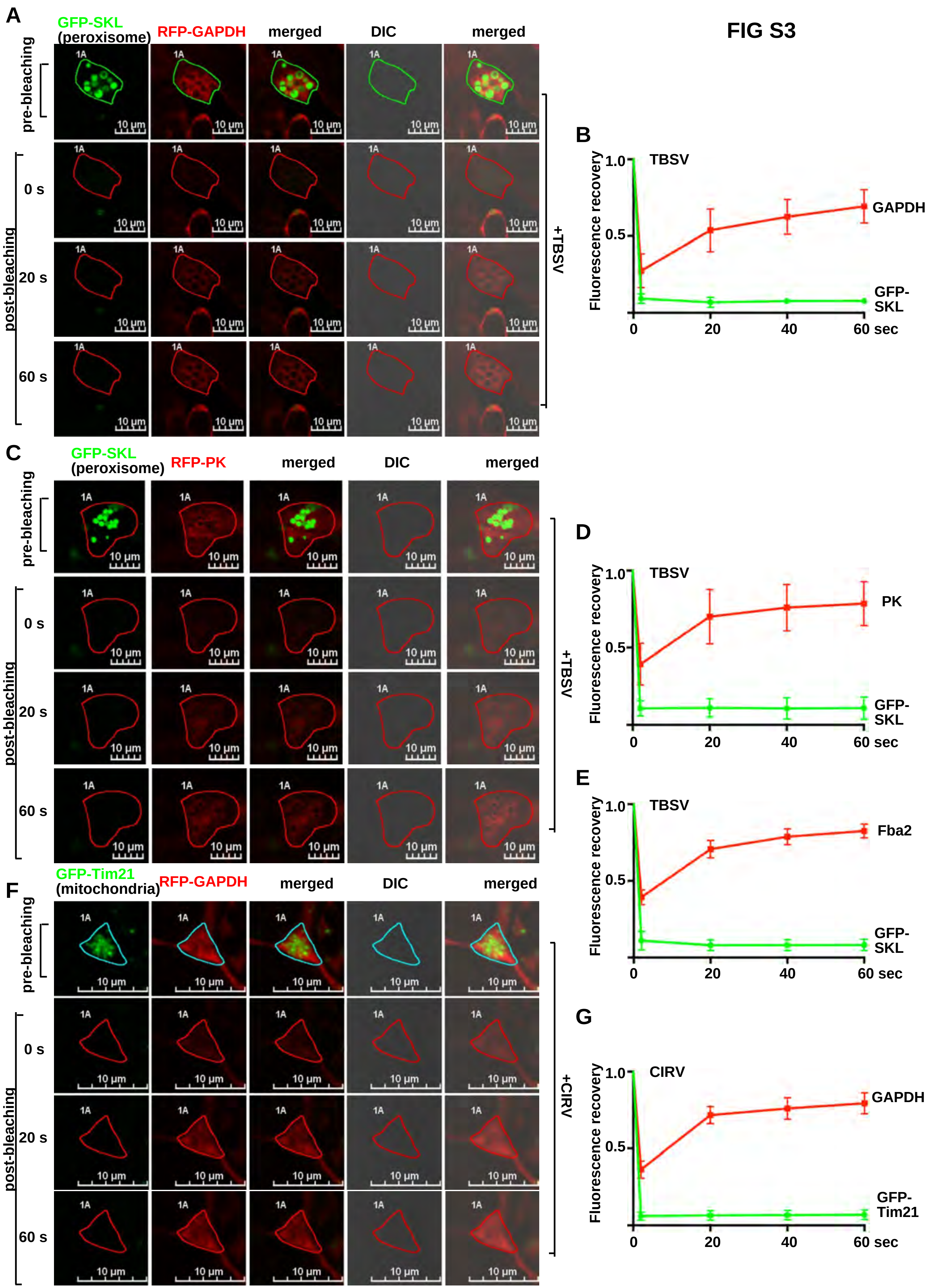

A
GFP-SKL
(peroxisome)
RFP-GAPDH
merged
DIC
merged
pre-bleaching
post-bleaching
0 s
20 s
60 s
1A
10 µm
+TBSV
B
TBSV
Fluorescence recovery
1.0
0.5
GAPDH
GFP-SKL
0
20
40
60 sec
C
GFP-SKL
(peroxisome)
RFP-PK
merged
DIC
merged
D
TBSV
PK
GFP-SKL
E
TBSV
Fba2
GFP-SKL
F
GFP-Tim21
(mitochondria)
RFP-GAPDH
merged
DIC
merged
+CIRV
G
CIRV
GAPDH
GFP-Tim21

FIG S4

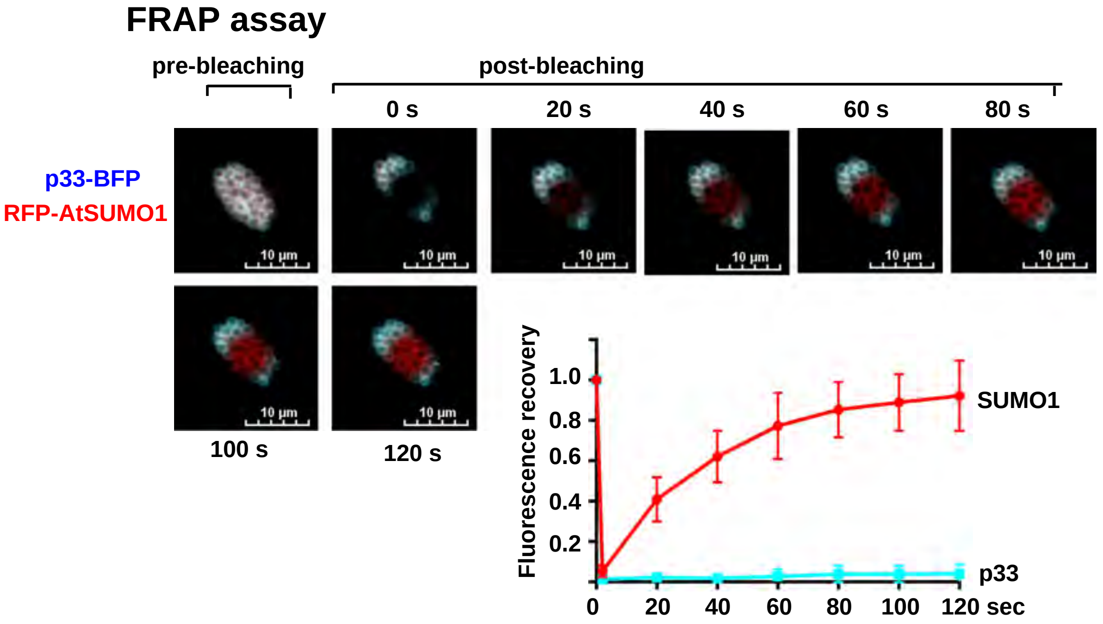
FRAP assay
pre-bleaching
post-bleaching
0 s
20 s
40 s
60 s
80 s
p33-BFP
RFP-AtSUMO1
10 µm
100 s
120 s
Fluorescence recovery
1.0
0.8
0.6
0.4
0.2
0
20
40
60
80
100
120 sec
SUMO1
p33

#### FIG S5

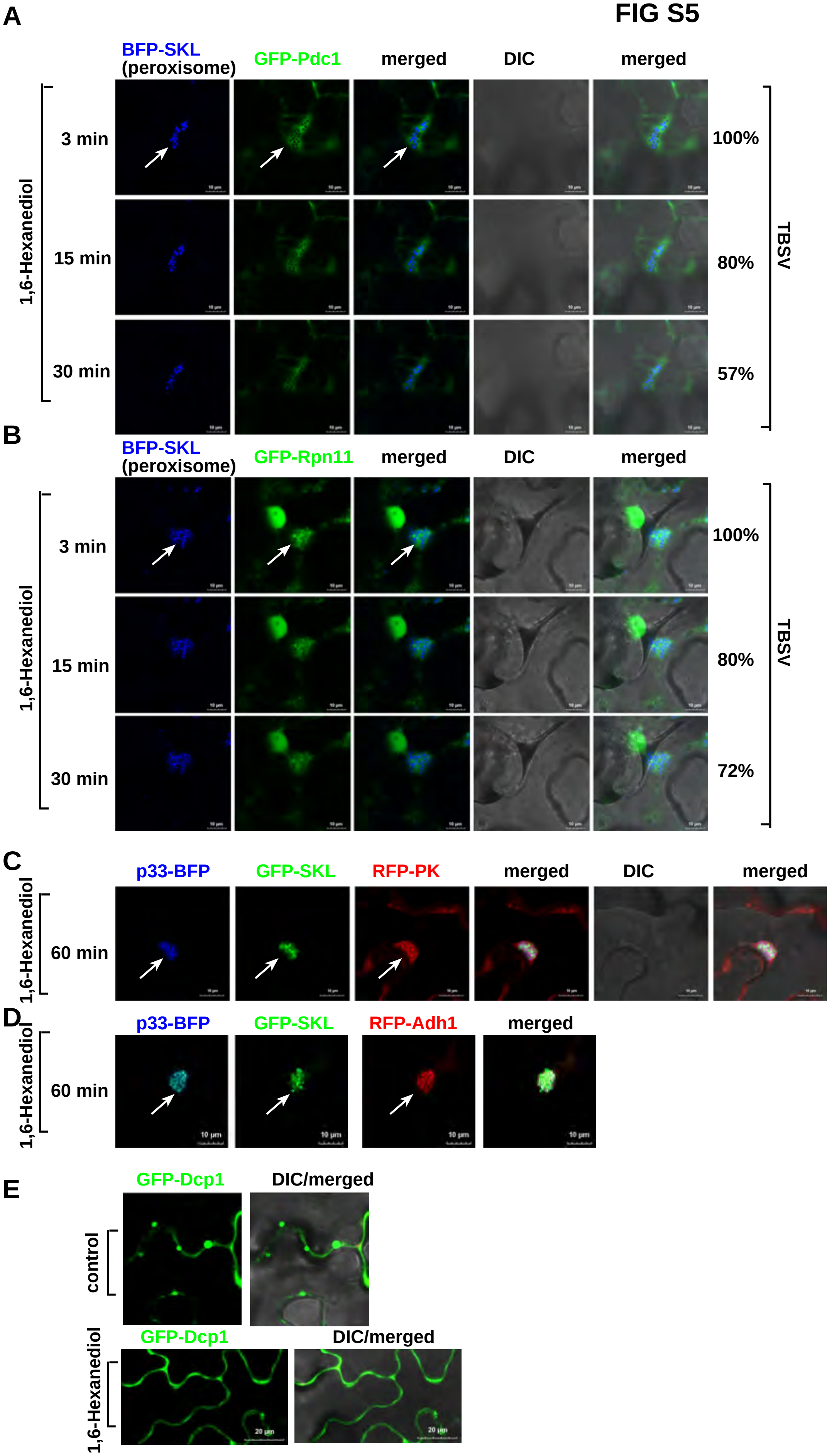

A
BFP-SKL (peroxisome)
GFP-Pdc1
merged
DIC
merged
1,6-Hexanediol
3 min
15 min
30 min
100%
80%
57%
TBSV
B
BFP-SKL (peroxisome)
GFP-Rpn11
merged
DIC
merged
1,6-Hexanediol
3 min
15 min
30 min
100%
80%
72%
TBSV
C
p33-BFP
GFP-SKL
RFP-PK
merged
DIC
merged
1,6-Hexanediol
60 min
D
p33-BFP
GFP-SKL
RFP-Adh1
merged
1,6-Hexanediol
60 min
10 µm
E
GFP-Dcp1
DIC/merged
control
1,6-Hexanediol
20 µm

FIG S6

#### droplet formation assay

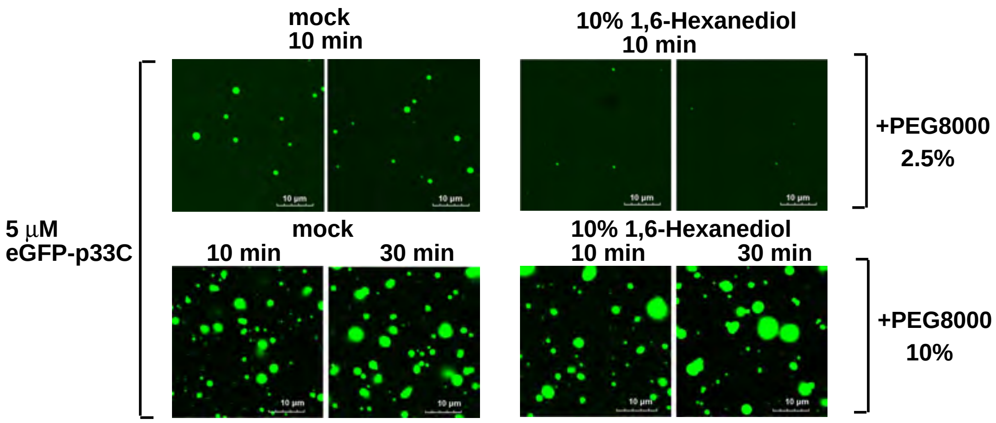

FIG S7

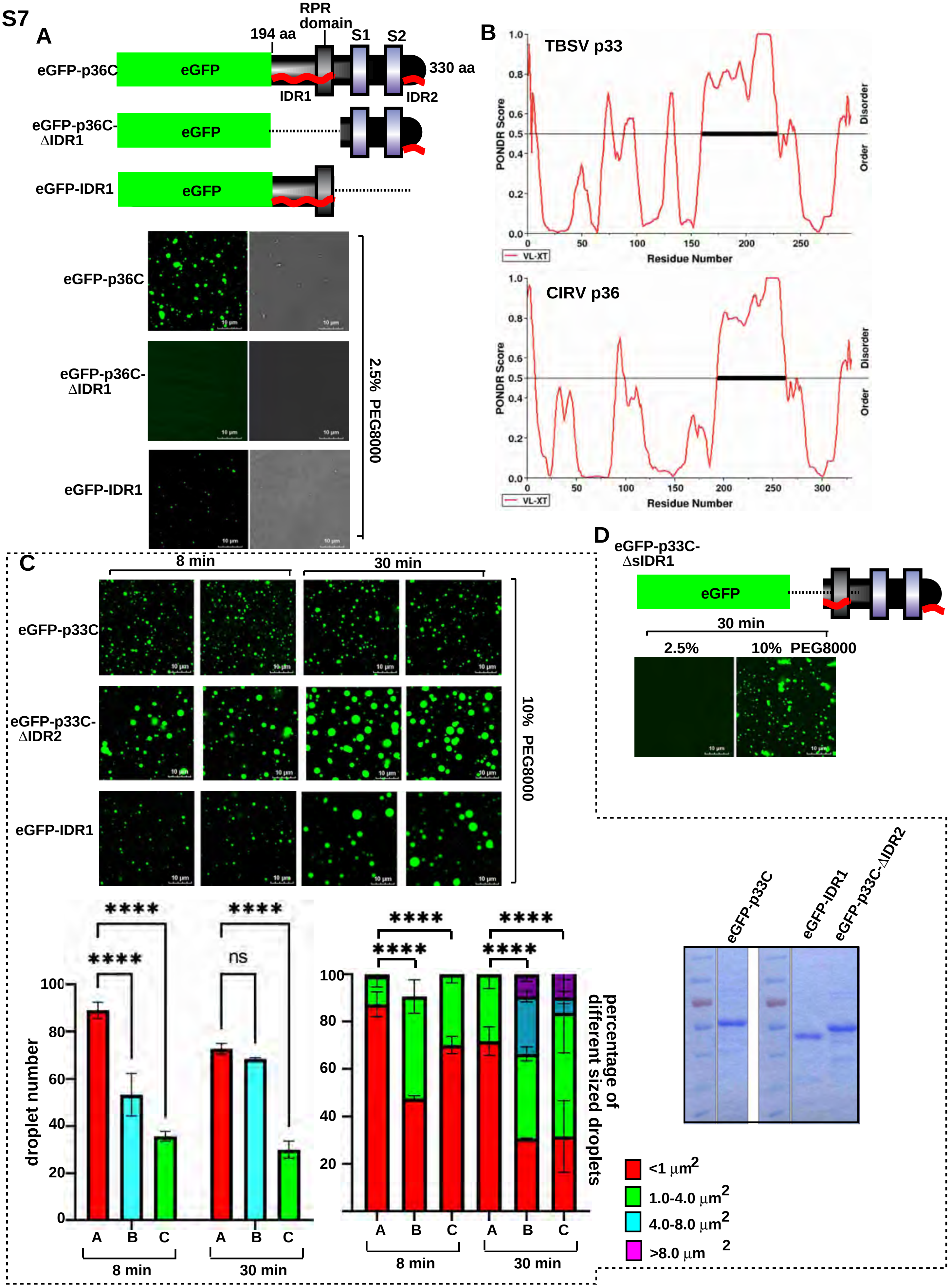

A
194 aa
RPR domain
S1
S2
eGFP-p36C
eGFP
330 aa
IDR1
IDR2
eGFP-p36C-ΔIDR1
eGFP-IDR1
2.5% PEG8000
B
TBSV p33
CIRV p36
PONDR Score
Disorder
Order
Residue Number
VL-XT
C
8 min
30 min
eGFP-p33C
eGFP-p33C-ΔIDR2
10% PEG8000
droplet number
ns
A
B
C
percentage of different sized droplets
<1 μm2
1.0-4.0 μm2
4.0-8.0 μm2
>8.0 μm2
D
eGFP-p33C-ΔsIDR1
2.5%
10% PEG8000
eGFP-p33C
eGFP-IDR1
eGFP-p33C-ΔIDR2

### FIG S8

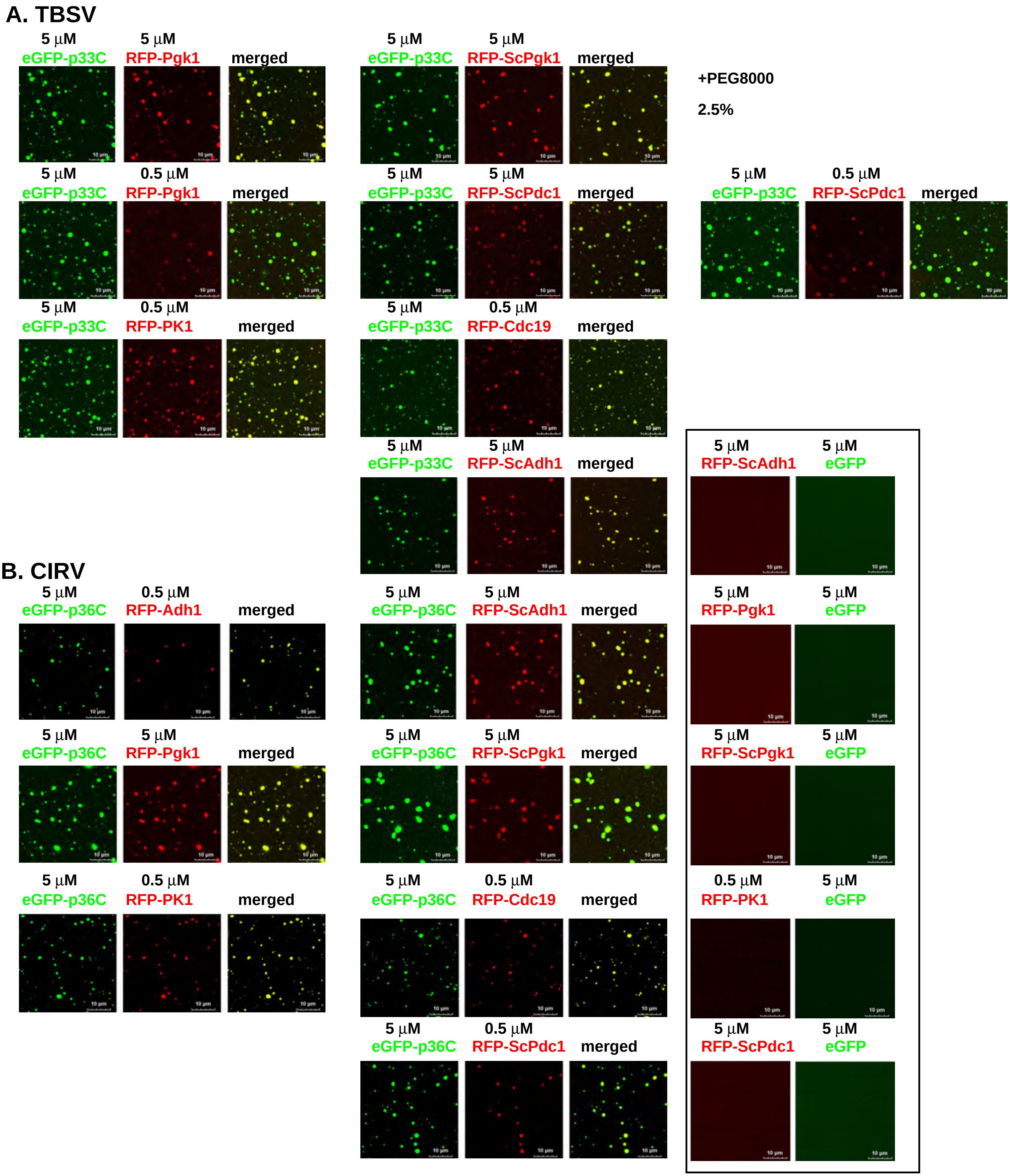
A. TBSV
5 µM eGFP-p33C
5 µM RFP-Pgk1
merged
5 µM eGFP-p33C
5 µM RFP-ScPgk1
merged
+PEG8000
2.5%
5 µM eGFP-p33C
0.5 µM RFP-Pgk1
merged
5 µM eGFP-p33C
5 µM RFP-ScPdc1
merged
5 µM eGFP-p33C
0.5 µM RFP-ScPdc1
merged
5 µM eGFP-p33C
0.5 µM RFP-PK1
merged
5 µM eGFP-p33C
0.5 µM RFP-Cdc19
merged
5 µM eGFP-p33C
5 µM RFP-ScAdh1
merged
5 µM RFP-ScAdh1
5 µM eGFP
B. CIRV
5 µM eGFP-p36C
0.5 µM RFP-Adh1
merged
5 µM eGFP-p36C
5 µM RFP-ScAdh1
merged
5 µM RFP-Pgk1
5 µM eGFP
5 µM eGFP-p36C
5 µM RFP-Pgk1
merged
5 µM eGFP-p36C
5 µM RFP-ScPgk1
merged
5 µM RFP-ScPgk1
5 µM eGFP
5 µM eGFP-p36C
0.5 µM RFP-PK1
merged
5 µM eGFP-p36C
0.5 µM RFP-Cdc19
merged
0.5 µM RFP-PK1
5 µM eGFP
5 µM eGFP-p36C
0.5 µM RFP-ScPdc1
merged
5 µM RFP-ScPdc1
5 µM eGFP

FIG S9

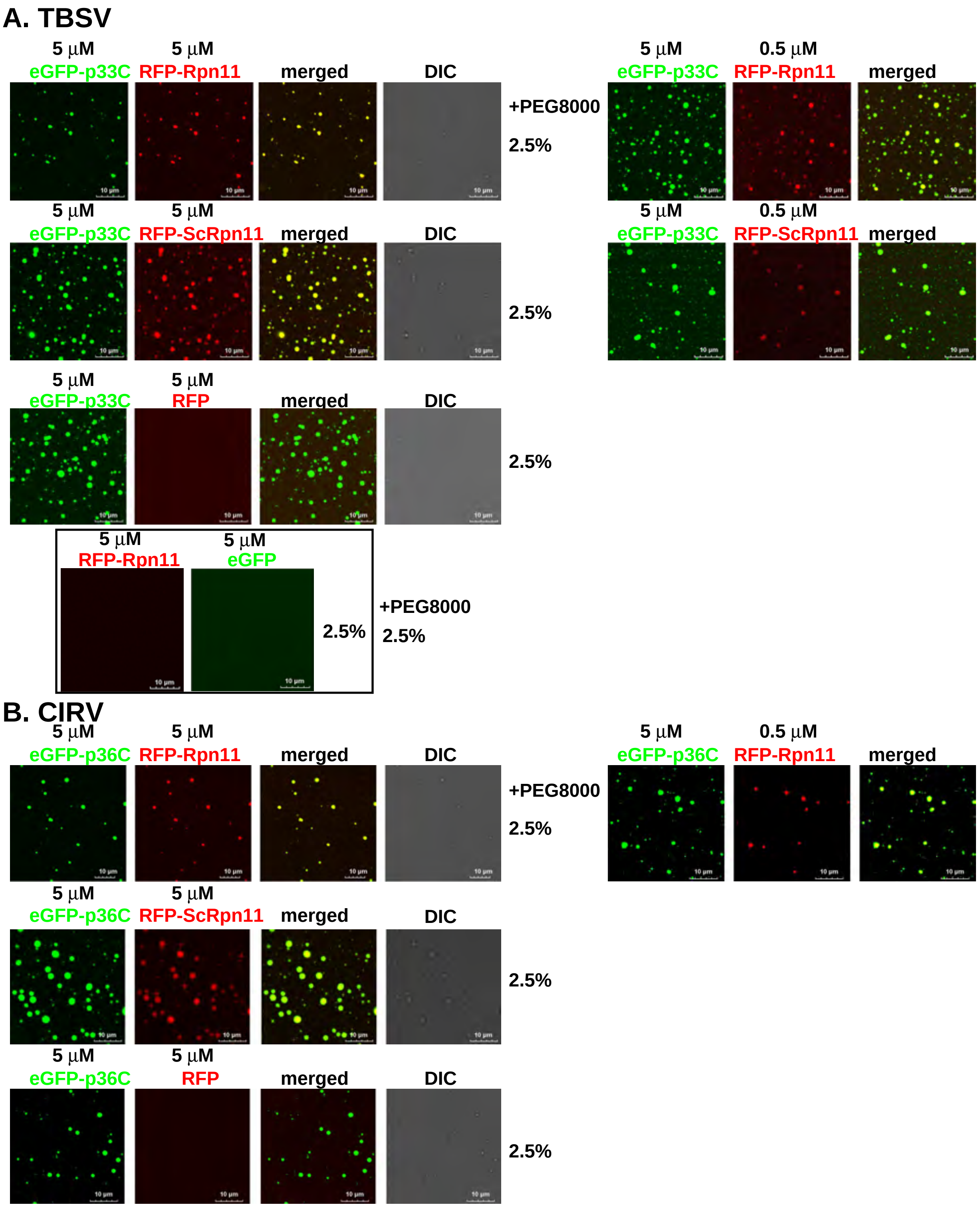

A. TBSV
5 µM
eGFP-p33C
5 µM
RFP-Rpn11
merged
DIC
+PEG8000
2.5%
5 µM
eGFP-p33C
0.5 µM
RFP-Rpn11
merged
5 µM
eGFP-p33C
5 µM
RFP-ScRpn11
merged
DIC
2.5%
5 µM
eGFP-p33C
0.5 µM
RFP-ScRpn11
merged
5 µM
eGFP-p33C
5 µM
RFP
merged
DIC
2.5%
5 µM
RFP-Rpn11
5 µM
eGFP
2.5%
+PEG8000
2.5%
B. CIRV
5 µM
eGFP-p36C
5 µM
RFP-Rpn11
merged
DIC
+PEG8000
2.5%
5 µM
eGFP-p36C
0.5 µM
RFP-Rpn11
merged
5 µM
eGFP-p36C
5 µM
RFP-ScRpn11
merged
DIC
2.5%
5 µM
eGFP-p36C
5 µM
RFP
merged
DIC
2.5%
10 µm

#### FIG S10

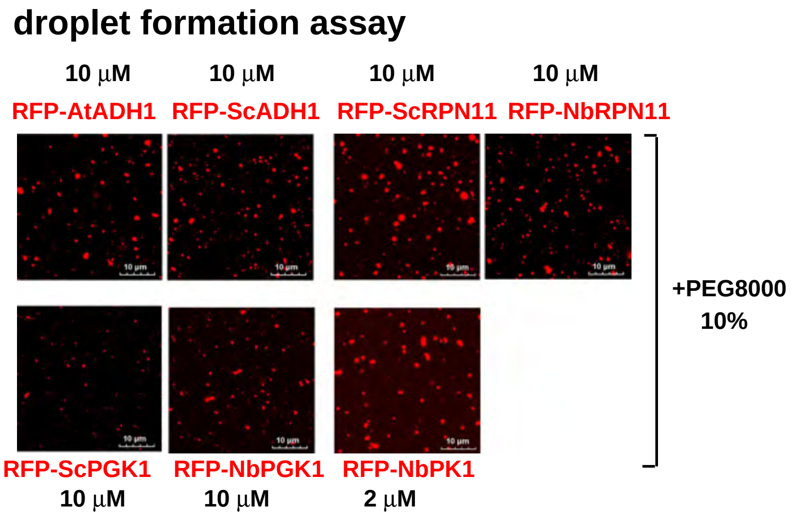
droplet formation assay
10 µM
RFP-AtADH1
10 µM
RFP-ScADH1
10 µM
RFP-ScRPN11
10 µM
RFP-NbRPN11
+PEG8000
10%
RFP-ScPGK1
10 µM
RFP-NbPGK1
10 µM
RFP-NbPK1
2 µM

#### FIG S11

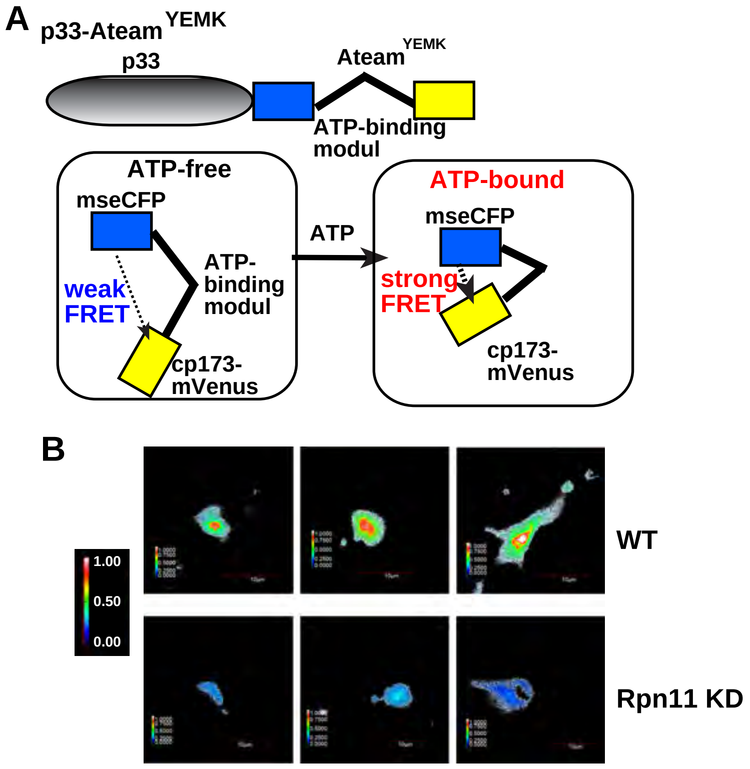
A
p33-Ateam YEMK
p33
Ateam YEMK
ATP-binding modul
ATP-free
mseCFP
ATP-binding modul
weak FRET
cp173-mVenus
ATP
ATP-bound
mseCFP
strong FRET
cp173-mVenus
B
1.00
0.50
0.00
WT
Rpn11 KD

FIG S12

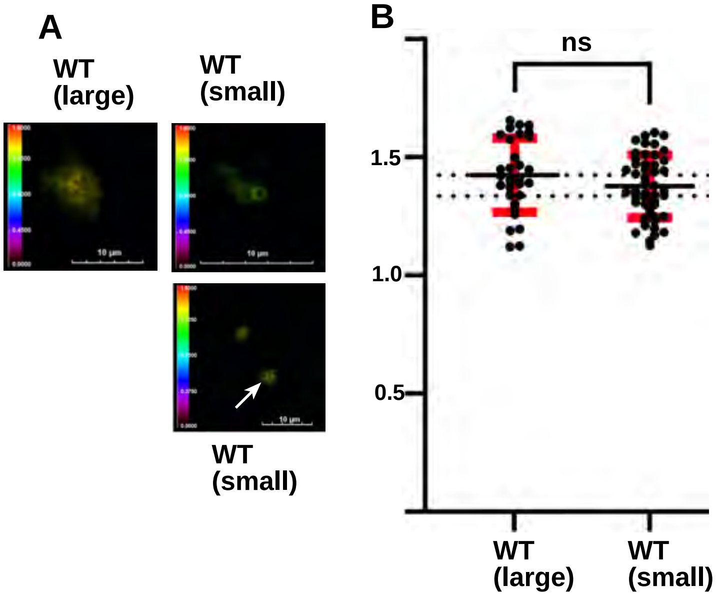
A
WT (large)
WT (small)
10 µm
10 µm
10 µm
WT (small)
B
ns
1.5
1.0
0.5
WT (large)
WT (small)

FIG S13

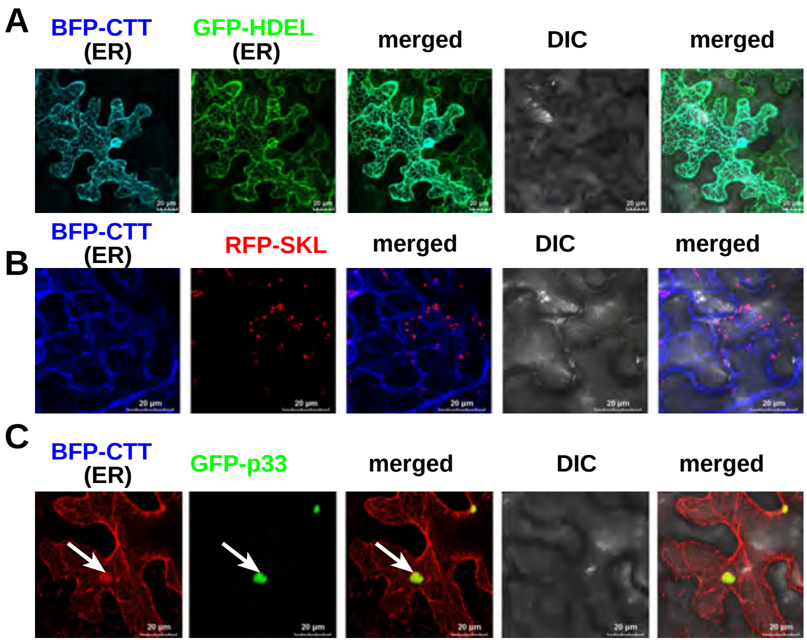
A
BFP-CTT
(ER)
GFP-HDEL
(ER)
merged
DIC
merged
B
BFP-CTT
(ER)
RFP-SKL
merged
DIC
merged
C
BFP-CTT
(ER)
GFP-p33
merged
DIC
merged

FIG S14

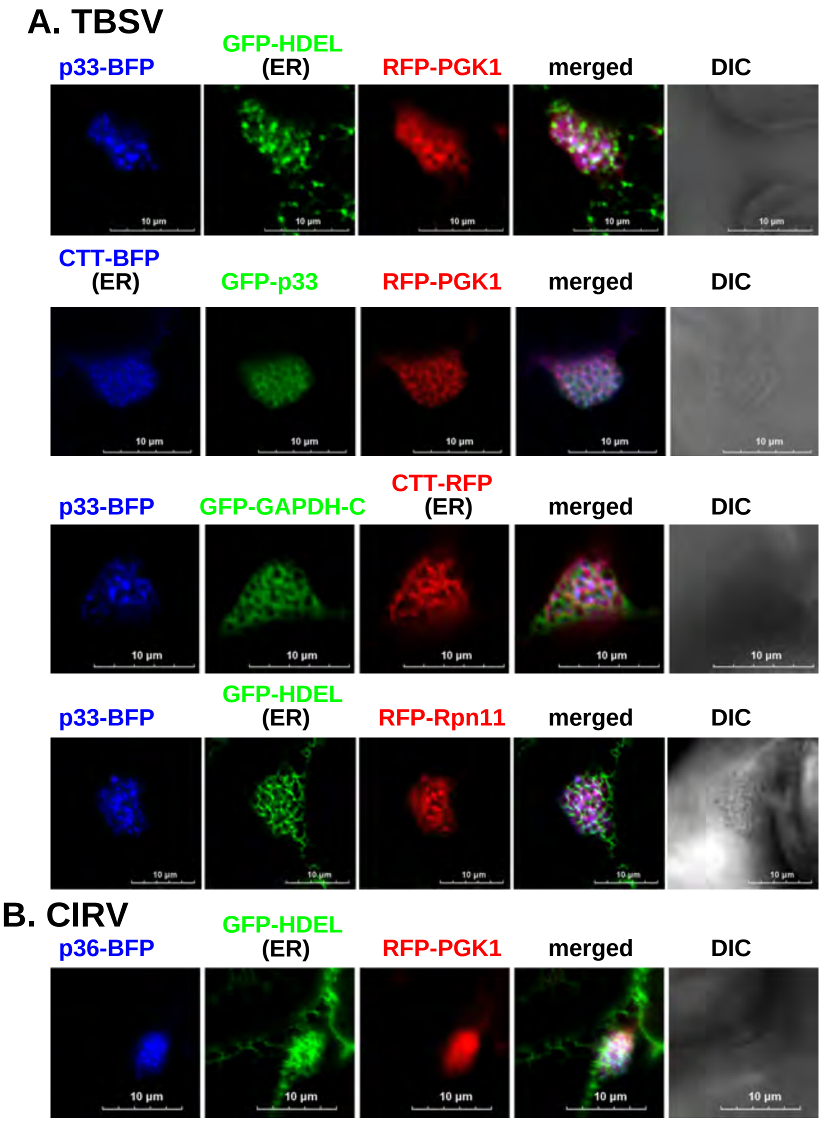

A. TBSV
p33-BFP
GFP-HDEL (ER)
RFP-PGK1
merged
DIC
CTT-BFP (ER)
GFP-p33
RFP-PGK1
merged
DIC
p33-BFP
GFP-GAPDH-C
CTT-RFP (ER)
merged
DIC
p33-BFP
GFP-HDEL (ER)
RFP-Rpn11
merged
DIC
B. CIRV
p36-BFP
GFP-HDEL (ER)
RFP-PGK1
merged
DIC
10 µm

FIG S15

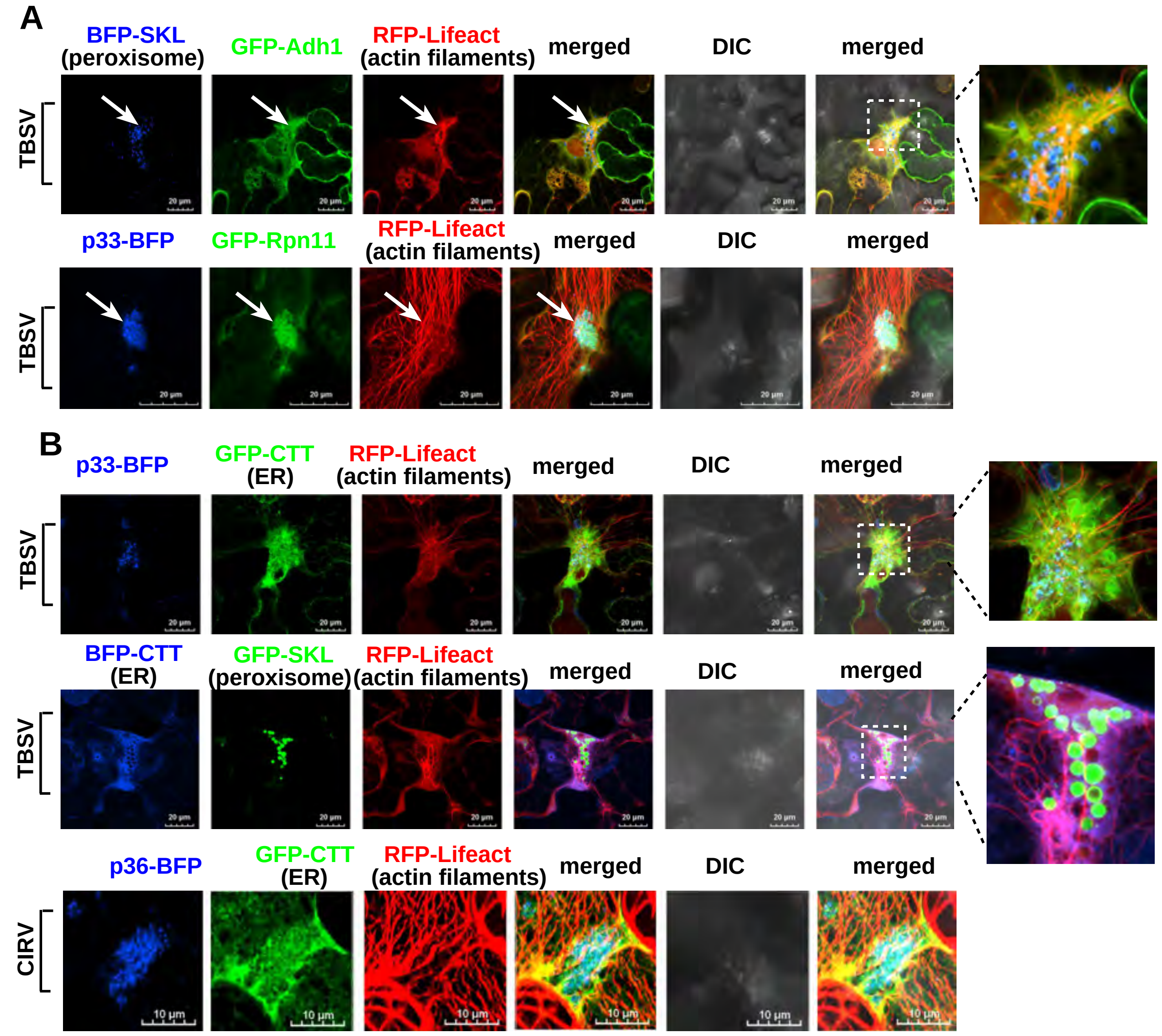

A
BFP-SKL
(peroxisome)
GFP-Adh1
RFP-Lifeact
(actin filaments)
merged
DIC
merged
TBSV
p33-BFP
GFP-Rpn11
RFP-Lifeact
(actin filaments)
merged
DIC
merged
TBSV
B
p33-BFP
GFP-CTT
(ER)
RFP-Lifeact
(actin filaments)
merged
DIC
merged
TBSV
BFP-CTT
(ER)
GFP-SKL
(peroxisome)
RFP-Lifeact
(actin filaments)
merged
DIC
merged
TBSV
p36-BFP
GFP-CTT
(ER)
RFP-Lifeact
(actin filaments)
merged
DIC
merged
CIRV
10 µm

#### FIG S16

##### BiFC assay

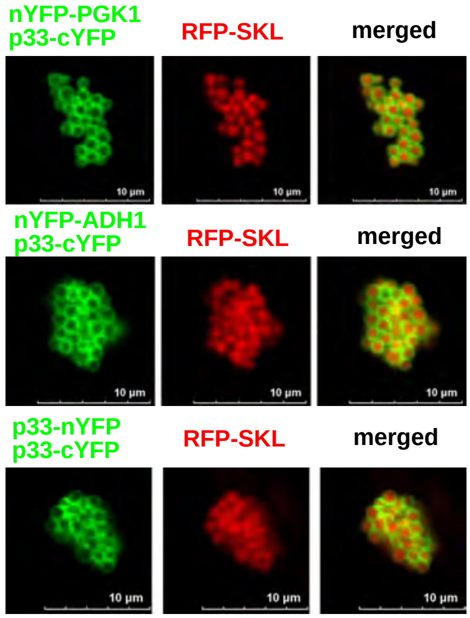

### Plasmids constructed in this study

| Number | Plasmid name | insert source | insert digestion sites | primers used for insert amplification | vector source | vector digestion sites |
|---|---|---|---|---|---|---|
| 1 | pGD-RFP-NbPK | N.benthamiana cDNA | BglII/SalI | 6659/6660 | pGD-RFP | BamHI/SalI |
| 2 | pRS315-Tef-mRFP-ScCdc19 | yeast genomic DNA | BamHI/XhoI | 5992/5993 | LpRS315-Tef-mRFP-MCS | BamHI/SalI |
| 3 | pRS315-Tef-mRFP-ScPgk1 | yeast genomic DNA | BclI/XhoI | 8599/6344 | LpRS315-Tef-mRFP-MCS | BamHI/SalI |
| 4 | pRS315-Tef-mRFP-ScPdc1 | yeast genomic DNA | BamHI/XhoI | 5621/2409 | LpRS315-Tef-mRFP-MCS | BamHI/SalI |
| 5 | pRS315-Tef-mRFP-ScAdh1 | yeast genomic DNA | BamHI/XhoI | 7574/7575 | LpRS315-Tef-mRFP-MCS | BamHI/SalI |
| 6 | pRS315-Tef-mRFP-ScRpn11 | yeast genomic DNA | BamHI/XhoI | 5234/5448 | LpRS315-Tef-mRFP-MCS | BamHI/SalI |
| 7 | pGD-GFP-AtGAPDH-C | Arabidposis thaliana cDNA | BamHI/BglII | 2067/8705 | pGD-eGFP | BamHI |
| 8 | pGD-RFP-AtGAPDH-C | Arabidposis thaliana cDNA | BamHI/BglII | 2067/8705 | pGD-RFP | BamHI |
| 9 | pGD-TagBFP-SKL | pGD-N-tagBFP | BglII/XhoI | 5673/5676 | pGD-35S | BamHI/XhoI |
| 10 | pET-30a-RFP | pGD-N-RFP | BglII/XhoI | 5649/5652 | pET-30a | BamHI/XhoI |
| 11 | pET-30a-mRFP-NbPgk1 | pGD-RFP-NbPGK | BglII/SalI | 5649/6439 | pET-30a | BamHI/XhoI |
| 12 | pET-30a-mRFP-AtAdh1 | pGD-RFP-AtAdh1 | BglII/XhoI | 5649/7908 | pET-30a | BamHI/XhoI |
| 13 | pET-30a-mRFP-NbPK | pGD-RFP-NbPK | BglII/SalI | 5649/6660 | pET-30a | BamHI/XhoI |
| 14 | pET-30a-mRFP-ScCdc19 | pRS315-Tef-mRFP-ScCdc19 | SacI/XhoI | 8725/5993 | pET-30a | SacI/XhoI |
| 15 | pET-30a-mRFP-ScPgk1 | pRS315-Tef-mRFP-ScPgk1 | BamHI/XhoI | 2630/6344 | pET-30a | BamHI/XhoI |
| 16 | pET-30a-mRFP-ScPdc1 | pRS315-Tef-mRFP-ScPdc1 | SacI/XhoI | 8725/2409 | pET-30a | SacI/XhoI |
| 17 | pET-30a-mRFP-ScAdh1 | pRS315-Tef-mRFP-ScAdh1 | SacI/XhoI | 8725/7575 | pET-30a | SacI/XhoI |
| 18 | pET-30a-mRFP-ScRpn11 | pRS315-Tef-mRFP-ScRpn11 | SacI/XhoI | 8725/5448 | pET-30a | SacI/XhoI |
| 19 | pGD-BFP-NbADH1 | N.benthamiana cDNA | BamHI/XhoI | 8726/8727 | pGD-BFP | BamHI/SalI |
| 20 | pGD-RFP-NbADH1 | N.benthamiana cDNA | BamHI/XhoI | 8726/8727 | pGD-RFP | BamHI/SalI |
| 21 | pGD-RFP-NbPdc1 | N.benthamiana cDNA | BamHI/XhoI | 8732/8733 | pGD-RFP | BamHI/SalI |
| 22 | pGD-BFP-NbRpn11 | N.benthamiana cDNA | BamHI/XhoI | 8730/8731 | pGD-BFP | BamHI/SalI |
| 23 | pGD-RFP-NbRpn11 | N.benthamiana cDNA | BamHI/XhoI | 8730/8731 | pGD-RFP | BamHI/SalI |
| 24 | pET-30a-eGFP | pGD-eGFP | BglII/XhoI | 6532/2599 | pET-30a | BamHI/XhoI |

| | | | | | | |
|---|---|---|---|---|---|---|
| 25 | pRS315-Tef-mRFP-NbRpn11 | N.benthamiana cDNA | BamHI/XhoI | 8730/8731 | pRS315-Tef-mRFP | BamHI/SalI |
| 26 | pET-30a-mRFP-NbRpn11 | pRS315-Tef-mRFP-NbRpn11 | SacI/XhoI | 8725/8731 | pET-30a | SacI/XhoI |
| 27 | pGD-eGFP-T33c | pGD-T33-cYFP | BglII/XhoI | 6879/1593 | pGD-eGFP | BamHI/SalI |
| 28 | pET-30a-eGFP-T33c | pGD-eGFP-T33c | BglII/XhoI | 6532/1593 | pET-30a | BamHI/XhoI |
| 29 | pGD-eGFP-T33-△IDR1(△160-229aa) | pGD-T33-cYFP | BglII/XhoI | 4000/8756+8755/1593 | pGD-eGFP | BamHI/SalI |
| 30 | pGD-eGFP-C36-△IDR1(△194-262aa) | pGD-C36-cYFP | BamHI/XhoI | 6184/8759+8758/3665 | pGD-eGFP | BamHI/SalI |
| 31 | pGD-BFP-CTT-ER | Arabidposis thaliana cDNA | BamHI/XhoI | 8760/8761 | pGD-BFP | BamHI/SalI |
| 32 | pGD-eGFP-CTT-ER | Arabidposis thaliana cDNA | BamHI/XhoI | 8760/8761 | pGD-eGFP | BamHI/SalI |
| 33 | pGD-eGFP-C36-194-262aa(eGFP-C36-IDR1) | pGD-C36-cYFP | BamHI/XhoI | 8765/8766 | pGD-eGFP | BamHI/SalI |
| 34 | pGD-eGFP-C36-263-329aa(eGFP-C36c-△IDR1) | pGD-C36-cYFP | BamHI/XhoI | 8763/3665 | pGD-eGFP | BamHI/SalI |
| 35 | pGD-eGFP-C36 | pGD-C36-cYFP | BamHI/XhoI | 3461/3665 | pGD-eGFP | BamHI/SalI |
| 36 | pGD-eGFP-C36-194-329aa(eGFP-C36c) | pGD-C36-cYFP | BamHI/XhoI | 8765/3665 | pGD-eGFP | BamHI/SalI |
| 37 | pET-30a-eGFP-C36-194-262aa(eGFP-C36-IDR1) | pGD-eGFP-C36-194-262aa | BglII/XhoI | 6532/8766 | pET-30a | BamHI/XhoI |
| 38 | pET-30a-eGFP-C36-263-329aa(eGFP-C36c-△IDR1) | pGD-eGFP-C36-263-329aa | BglII/XhoI | 6532/3665 | pET-30a | BamHI/XhoI |
| 39 | pET-30a-eGFP-C36-194-329aa(eGFP-C36c) | pGD-eGFP-C36-194-329aa | BglII/XhoI | 6532/3665 | pET-30a | BamHI/XhoI |
| 40 | pGD-eGFP-T33-N240(IDR1) | pGD-T33-cYFP | BglII/XhoI | 4000/5715 | pGD-eGFP | BamHI/SalI |
| 41 | pGD-eGFP-T33-1-181aa(sIDR1) | pGD-T33-cYFP | BglII/XhoI | 4000/5713 | pGD-eGFP | BamHI/SalI |
| 42 | pGD-eGFP-T33-WT | pGD-T33-cYFP | BglII/XhoI | 4000/1593 | pGD-eGFP | BamHI/SalI |
| 43 | pGD-eGFP-T33c | pGD-T33-cYFP | BglII/XhoI | 6879/1593 | pGD-eGFP | BamHI/SalI |

| | | | | | | |
|---|---|---|---|---|---|---|
| 44 | pET-30a-eGFP-T33c-155-296aa(eGFP-T33c) | pGD-eGFP-T33c | BglII/XhoI | 6532/1593 | pET-30a | BamHI/XhoI |
| 45 | pET-30a-eGFP-T33c-155-240aa(eGFP-T33c-IDR1) | pGD-eGFP-T33c | BglII/XhoI | 6532/5715 | pET-30a | BamHI/XhoI |
| 46 | pET-30a-eGFP-T33c-155-284aa(eGFP-T33c-△IDR2) | pGD-eGFP-T33c | BglII/XhoI | 6532/8757 | pET-30a | BamHI/XhoI |
| 47 | pGD-eGFP-T33-181-296aa(eGFP-T33c-△sIDR1) | pGD-T33-cYFP | BglII/XhoI | 8808/1593 | pGD-eGFP | BamHI/SalI |
| 48 | pET-30a-eGFP-T33c-181-296aa(eGFP-T33c-△sIDR1) | pGD-eGFP-T33-181-296aa | BglII/XhoI | 6532/1593 | pET-30a | BamHI/XhoI |
| 49 | pGD-eGFP-T33-IDR-mut1 | pGD-T33-cYFP | BglII/XhoI | 4000/8854+8853/1593 | pGD-eGFP | BamHI/SalI |
| 50 | HpESC-Gal-T33-IDR-mut1/DI72 | pGD-eGFP-T33-IDR-mut1 | BglII/XhoI | 4000/1593 | HpESC-Gal-Hisp33/Gal-DI72 | BamHI/XhoI |
| 51 | HpESC-Gal-T33-WT/DI72 | pGD-T33-cYFP | BglII/XhoI | 4000/1593 | HpESC-Gal-Hisp33/Gal-DI72 | BamHI/XhoI |
| 52 | pGD-eGFP-T33-155-240-IDR-mut1 | pGD-eGFP-T33-IDR-mut1 | BglII/XhoI | 8857/5715 | pGD-eGFP | BamHI/SalI |
| 53 | pET-30a-eGFP-T33-155-240-IDR-mut1 | pGD-eGFP-T33-155-240-IDR-mut1 | BglII/XhoI | 6532/5715 | pET-30a | BamHI/XhoI |
| 54 | pGD-L-T33-YEMK | pGD-T33-cYFP | BglII/XhoI | 4000/8872 | pGD-L-C36-YEMK | BamHI/XhoI |
| 55 | pGD-L-T33-△IDR-YEMK | pGD-eGFP-T33-△IDR | BglII/XhoI | 4000/8872 | pGD-L-C36-YEMK | BamHI/XhoI |
| 56 | pGD-L-T33-IDR-mut1-YEMK | pGD-eGFP-T33-IDR-mut1 | BglII/XhoI | 4000/8872 | pGD-L-C36-YEMK | BamHI/XhoI |
| 57 | pGD-T33-N240-BFP | pGD-T33-cYFP | BglII/PstI | 4000/8891 | pGD-C-BFP | BamHI/PstI |

### Other plasmids used in this study

| Number | Plasmid name | reference or origin |
|---|---|---|
| 1 | UpYES-NT vector | [1]Lin et al., 2021, Plos pathogens |
| 2 | LpGAD-Gal-HisT92 | [1]Lin et al., 2021, Plos pathogens |
| 3 | pGD-2X 35s L vector | [2]Xu and Nagy, 2016, Plos biology&[3]Xu and Nagy, 2015, PNAS |

| | | |
|---|---|---|
| 4 | pGD-2X 35s L N-RFP | [2]Xu and Nagy, 2016, Plos biology&[3]Xu and Nagy, 2015, PNAS |
| 5 | pGD-2X 35s L N-BFP | [2]Xu and Nagy, 2016, Plos biology&[3]Xu and Nagy, 2015, PNAS |
| 6 | pGD-2X 35s L C-RFP | [2]Xu and Nagy, 2016, Plos biology&[3]Xu and Nagy, 2015, PNAS |
| 7 | pGD-2X 35s L C-BFP | [2]Xu and Nagy, 2016, Plos biology&[3]Xu and Nagy, 2015, PNAS |
| 8 | pGD-T33-RFP | [2]Xu and Nagy, 2016, Plos biology&[3]Xu and Nagy, 2015, PNAS |
| 9 | pGD-T33-BFP | [2]Xu and Nagy, 2016, Plos biology&[3]Xu and Nagy, 2015, PNAS |
| 10 | pGD-At Pex13-RFP | [2]Xu and Nagy, 2016, Plos biology&[3]Xu and Nagy, 2015, PNAS |
| 11 | pGD-rsGFP-SKL | [2]Xu and Nagy, 2016, Plos biology&[3]Xu and Nagy, 2015, PNAS |
| 18 | pGD-N-eGFP | [2]Xu and Nagy, 2016, Plos biology&[3]Xu and Nagy, 2015, PNAS |
| 19 | pRS315-N-mRFP | [2]Xu and Nagy, 2016, Plos biology&[3]Xu and Nagy, 2015, PNAS |
| 12 | pGD-RFP-SKL | [2]Xu and Nagy, 2016, Plos biology&[3]Xu and Nagy, 2015, PNAS |
| 13 | pGD-CIRV-P36-TeamYEMK | [3]Chuang et al., 2017,Cell host & microbe |
| 14 | pGD-GFP-AtRPN11 | [4]Molho et al., 2021, Plos pathogens |
| 15 | pGD-RFP-AtRpn11 | [4]Molho et al., 2021, Plos pathogens |
| 16 | pGD-RFP-AtFba2 | [4]Molho et al., 2021, Plos pathogens |
| 17 | pGD-RFP-NbPgk1 | [4]Molho et al., 2021, Plos pathogens |
| 20 | ER-gb-GFP-HDEL | [5]Inaba at al., 2019, PNAS |
| 21 | pGD-GFP-AtPdc1 | [6]Lin et al., 2019, Plos pathogens |
| 22 | pGD-GFP-AtAdh1 | [6]Lin et al., 2019, Plos pathogens |
| 23 | pGD-CoxIV-BFP | gift from Mr Biao Sun(University of Kentucky) |
| 24 | pGD-35S-DI72-sat | gift from Dr. Daniel Barajas(University of Kentucky) |
| 25 | pGD-35S-L-p92 | gift from Dr. Daniel Barajas(University of Kentucky) |
| 26 | pB7m24GW-pro35S-lifeact-mRuby3 | gift from Dr. Tomo Kawashima(University of Kentucky) |


[1]Lin W, Feng Z, Prasanth K R, et al. Dynamic interplay between the co-opted Fis1 mitochondrial fission protein and membrane contact site proteins in supporting tombusvirus replication[J]. PLoS Pathogens, 2021, 17(3): e1009423.

[2]Xu K, Nagy P D. Enrichment of phosphatidylethanolamine in viral replication compartments via co-opting the endosomal Rab5 small GTPase by a positive-strand RNA virus[J]. PLoS Biology, 2016, 14(10): e2000128.

[3]Xu K, Nagy P D. RNA virus replication depends on enrichment of phosphatidylethanolamine at replication sites in subcellular membranes[J]. Proceedings of the National Academy of Sciences, 2015, 112(14): E1782-E1791.

[4]Chuang C, Prasanth K R, Nagy P D. The glycolytic pyruvate kinase is recruited directly into the viral replicase complex to generate ATP for RNA synthesis[J]. Cell host & microbe, 2017, 22(5): 639-652. e7.

[5]Molho M, Lin W, Nagy P D. A novel viral strategy for host factor recruitment: The co-opted proteasomal Rpn11 protein interaction hub in cooperation with subverted actin filaments are targeted to deliver cytosolic host factors for viral replication[J]. PLoS pathogens, 2021, 17(6): e1009680.

[6]Inaba J, Xu K, Kovalev N, et al. Screening Legionella effectors for antiviral effects reveals Rab1 GTPase as a proviral factor coopted for tombusvirus replication[J]. Proceedings of the National Academy of Sciences, 2019, 116(43): 21739-21747.

[7]Lin W, Liu Y, Molho M, et al. Co-opting the fermentation pathway for tombusvirus replication: Compartmentalization of cellular metabolic pathways for rapid ATP generation[J]. PLoS Pathogens, 2019, 15(10): e1008092.

### Primers used in this study

| Number | Oligo number | Oligo name | Oligo sequence |
|---|---|---|---|
| 1 | 1593 | 1593/TBSV/33stop/Xho/R | cggCTCGAGctatttgacacccagggactcctgt |
| 2 | 2067 | 2067/atGAPDH/Bam/F | cgagggatccatggctgacaagaagattaggatcggaatc |
| 3 | 2409 | 2409/PDC1/Xho/R | CgcgCTCGAGTTATTGCTTAGCGTTGGTAGC |
| 4 | 2599 | 2599/EYFP/XhoI/R | CggcCTCGAGTTACTTGTACAGCTCGTCCATGCCGA |
| 5 | 2630 | 2630/RFPmono/Bam/F | CgcgGGATCCatggcctcctccgaggacgtc |
| 6 | 3461 | 3461/CIRVp36/BamHI(2)/F | GccGGATCCatggagggtttgaaggctg |
| 7 | 3665 | 3665/CIRVp36/stop/XhoI/R | cgcgCTCGAGctatttgacaccgagggattc |
| 8 | 4000 | 4000/TBSV33/Bgl/F | ccagagatctatggagaccatcaagagaatg |
| 9 | 5234 | 5234/RPN11/BamHI/F | CGCGGATCCATGGAACGACTACAGAGATT |
| 10 | 5448 | 5448/RPN11/XhoI/R | CCGCTCGAGTTATTTAATTGCCACTGAATT |
| 11 | 5621 | 5621/ScPDC1/BamHI/F | cgccggatccATGTCTGAAATTACTTTGGGTAAATA |
| 12 | 5649 | 5649/tagRFP/BglII/F | ggaagatctATGGTGTCTAAGGGCGAAGAG |
| 13 | 5652 | 5652/tagRFP/XhoI /R | ccgctcgagTTAATTAAGTTTGTGCCCCAGTTTGC |
| 14 | 5673 | 5673/tagBFP/BglII/F | ggaagatctATGTCTGAATTGATTAAAGAGAATATGC |
| 15 | 5676 | 5676/tagBFP/SKL/XhoI/R | ccgctcgagTTACAACTTAGAaccgccaccGTTCAATTTGTGTCCTAACTTAGAAGG |
| 16 | 5713 | 5713/TBSVp33/181aa/Xho/R | ccgctcgagTTAACAATCTGTCGCTTCTCTCTC |
| 17 | 5715 | 5715/TBSVp33/240aa/Xho/R | ccgctcgagTTAATTCTCTGGACTGTTCTTAAGGTAAC |
| 18 | 5837 | 5837/TBSV/p33/nostop/PstI/R | GCCctgcagTTTGACACCCAGGGACTCCTG |

| | | | |
|---|---|---|---|
| 19 | 5986 | 5986/AtRPN11/XhoI/R1 | CCGCTCGAGCTAGAAGACAACAGTGTCGA |
| 20 | 5992 | 5992/ScCDC19/BamHI/F | cgccggatccATGTCTAGATTAGAAAGATTGA |
| 21 | 5993 | 5993/ScCDC19/XhoI/R | cgccctcgagTTAAACGGTAGAGACTTGCAAAG |
| 22 | 6184 | 6184/CIRVp36/BamHI/Kozak/F | cgcggatcctaaacaATGGAGGGTTTGAAGGCTG |
| 23 | 6344 | 6344/PGK1/XhoI/R | CCGCTCGAGTTATTTCTTTTCGGATAAGAA |
| 24 | 6439 | 6439/PGK1-His (P)/SalI-R | CGCGTCGACTCAATGGTGATGGTGATGATGGGCATCATCTAGGGCAAGCA |
| 25 | 6532 | 6532/EGFP/BglII/F | ggaagatctATGGTGAGCAAGGGCGAG |
| 26 | 6659 | 6659/PK-P/BamHI-F | CGGGATCCATGGCGATTGAGAATAACAA |
| 27 | 6660 | 6660/PK-P/SalI-R | CGCGTCGACTCACTTGACTGTAACAATCTT |
| 28 | 6879 | 6879/TBSVp33/155aa/BglII/F | ggaagatctATGCCTAGGGAAAAACTGTCGGTATTTAAG |
| 29 | 7574 | 7574/ScADH1/BamHI/F | ACGCGGATCCATGTCTATCCCAGAAACTCAAAAAGG |
| 30 | 7575 | 7575/ScADH1/XhoI/R | ACCGCTCGAGTTATTTAGAAGTGTCAACAACGTATCTACC |
| 31 | 7908 | 7908/AtADH1/stop/XhoI/R | ACCGCTCGAGTCAAGCACCCATGGTGATGATGC |
| 32 | 8599 | 8599/ScPGK1-BclI/F | ACGCTGATCAATGTCTTTATCTTCAAAGTTGTCTGTC |
| 33 | 8705 | 8705/AtGAPDH-C-BglII-TGA-BamHI/R | ACGCGGATCCTCAAGATCTggcctttgacatgtggacgatcaag |
| 34 | 8725 | 8725/RFP-mono-SacI/F | CGCCGAGCTCatggcctcctccgaggacgtc |
| 35 | 8726 | 8726/NbAdh1-BamHI/F | ACGCGGATCCATGTCAACCAATACTGCTGGTC |
| 36 | 8727 | 8727/NbAdh1-XhoI/R | ACCGCTCGAGTCAATGTCCCATGGTGATCA |
| 37 | 8730 | 8730/NbRpn11-BamHI/F | ACGCGGATCCATGTCTGGAATGGAAAGATTGC |
| 38 | 8731 | 8731/NbRpn11-XhoI/R | ACCGCTCGAGTCAGAAGTCAACTGTGTCAAGC |
| 39 | 8732 | 8732/NbPdc1-BamHI/F | ACGCGGATCCATGGACGCGAAGATCAGAGC |
| 40 | 8733 | 8733/NbPdc1-XhoI/R | ACCGCTCGAGTTACTGAGGATTCGGTGGACGACTATTAG |
| 41 | 8755 | 8755/TBSVp33-delta160-229-OV/F | Ccctacctagggaaaaactgaaggtgggttaccttaagaac |
| 42 | 8756 | 8756/TBSVp33-delta160-229-OV/R | gttcttaaggtaacccaccttcagtttttccctaggtaggg |
| 43 | 8757 | 8757/T33/C36-deltaC12aa-XhoI-R | ACCGCTCGAGCTACGCCGACTCCTCCACTCC |
| 44 | 8758 | 8758/CIRVp36-delta194-262-OV-F | ctccccagggaaaaactgaaggtgggttaccttaagaatag |

| | | | |
|---|---|---|---|
| 45 | 8759 | 8758/CIRVp36-delta194-262-OV-F | ctccccagggaaaaactgaaggtgggttaccttaagaatag |
| 46 | 8760 | 8760/CTT-ER-BamHI/F | ACGCGGATCCATGgacaagacctctgaattcataatc |
| 47 | 8761 | 8761/CTT-ER-XhoI/R | ACCGCTCGAGCTACCCTGATTTGGTGTAGATACG |
| 48 | 8763 | 8763/T33-N230aa/C36-N263aa-BamHI/F | ACGCGGATCCATGAAGGTGGGTTACCTTAAGAA |
| 49 | 8765 | 8765/CIRVp36-N194aa-BamHI/F | ACGCGGATCCATGTCTGTGTTTAGGTTGAAATCAGAGG |
| 50 | 8766 | 8766/CIRVp36-N262aa-XhoI/R | ACCGCTCGAGCTAGGCTCTCGCGACCTGTG |
| 51 | 8806 | 8806/T33-Δ156-180-ovlp-1R | ccggctcaaccaccagacaaggtagggtagcgtacacgg |
| 52 | 8807 | 8807/T33-Δ156-180-ovlp-2F | ccgtgtacgctaccctaccttgtctggtggttgagccgg |
| 53 | 8808 | 8808/T33-181aa-BglII/F | ggaagatctATGtgtctggtggttgagccg |
| 54 | 8853 | 8853/T33-IDR-Mut1-ov/F | gcgggagcactgtcggtatttgggctgggaactgcggcaggagcacacatggaggatgag |
| 55 | 8854 | 8854/T33-IDR-Mut1-ov/R | Tcctgccgcagttcccagcccaaataccgacagtgctcccgcaggtagggtagcgtacac |
| 56 | 8857 | 8857/T33-155aa-Mut1-BglII/F | ggaagatctATGCCTGCGGGAGCACTGTCGG |
| 57 | 8872 | 8872/T33-XhoI-nonstop-R | ACCGCTCGAGTTTGACACCCAGGGACTCC |
| 58 | 8873 | 8873/T33-N181-XhoI-nonstop-R | ACCGCTCGAGACAATCTGTCGCTTCTCTCTC |
| 59 | 8891 | 8891/T33-N240-Xho-Pst-nonstop-R | ccgctcgagctgcagATTCTCTGGACTGTTCTTAAGGTAAC |
